# Enabling Greenhouse Gas Emission Reduction while Improving Rice Yield with a Methane-Derived Microbial Biostimulant

**DOI:** 10.1101/2024.03.13.584920

**Authors:** Sarma Rajeev Kumar, Einstein Mariya David, Gangigere Jagadish Pavithra, Gopalakrishnan Kumar Sajith, Kuppan Lesharadevi, Selvaraj Akshaya, Chavaddi Bassavaraddi, Gopal Navyashree, Panakanahalli Shivaramu Arpitha, Padmanabhan Sreedevi, Khan Zainuddin, Firdous Saiyyeda, Bondalakunta Ravindra Babu, Muralidhar Udagatti Prashanth, Ganesan Ravikumar, Palabhanvi Basavaraj, Chavana Sandeep Kumar, Vinod Munisanjeeviah Lakshmi Devi Kumar, Theivasigamani Parthasarathi, Ezhilkani Subbian

**Affiliations:** String Bio Private Limited, Vinayaka Nagar, Nagasandra Bangalore-560073, Karnataka, India; String Bio Private Limited, Centre for Cellular and Molecular Platforms, National Centre for Biological Sciences campus, Bangalore-560065, Karnataka, India; VIT School of Agricultural Innovations and Advanced Learning (VAIAL), Vellore Institute of Technology, Vellore-632014, Tamil Nadu, India; School of Biosciences and Technology (SBST), Vellore Institute of Technology, Vellore-632014, Tamil Nadu, India

**Keywords:** Climate Change, Methane, Microbial Biostimulant, Nitrous oxide, Global Warming Potential

## Abstract

Rice is a vital crop for food security and human nutrition, yet its cultivation produces ∼11% of total global anthropogenic methane (CH_4_) emissions - the second most important greenhouse gas (GHG). Modifications to rice management practice are necessary, both to increase yield and mitigate GHG emissions. We investigated the effect of a methane-derived microbial biostimulant on grain yield and GHG emissions from rice fields. Applications of microbial biostimulant resulted in significant enhancement of grain yield, even under different nitrogen management, with consistent reduction in GHG emissions. The study further outlines a potential mechanism for broad and diverse positive effects of microbial biostimulant on the paddy crop including in photosynthesis, tillering and panicle development. Observations from the study will help stakeholders and policy makers, leverage biological solutions like methane-derived microbial biostimulant to improve crop yield and address food security, while reducing anthropogenic CH4 emissions to meet targets agreed at COP26.

## 1. Introduction

Global climate change poses a significant threat to food security, presenting potentially existential economic, political, and social outcomes (Sova et al., 2019). Climate change negatively affects both food production and its quality. By 2050, the global population is projected to reach 10 billion, which will require a 70% increase in food production (van Dijk et al., 2021). For instance, by 2050, annual demand for cereals like maize, rice and wheat is projected to reach 3.3 billion tons or 800 million tons more than 2014’s combined harvest (FAO, 2016). For a food-secure future, global crop production will have to increase substantially and be climate resilient, while simultaneously reducing its environmental impact. Use of innovative technologies or approaches for achieving sustainable agriculture have been a matter of debate in the recent past. By proposing an ambitious agenda through the Paris Agreement and Sustainable Development Goals (SDGs), global leaders have acknowledged the urgent need to address climate change. New aggressive targets have been set in the COP26 meeting to reduce CH_4_ emissions and achieve net-zero by 2050 (Masood and Tollefson, 2021). However, even after several years of framing these policies, progress towards the targets is sobering.

Rice is one of the world’s top three staple crops and is closely connected with food security, economic growth, employment, culture, and regional peace. About 90% of the world’s rice is produced in Asia (FAO, 2019) and rice exports have been a key economic tool for this region. Rice paddies are also one of the most significant sources of CH_4_ and N_2_O emissions (Linquist et al., 2012; Carlson 2017; Timilsina et al., 2020; Qian et al., 2023). Global average annual CH_4_ emissions from rice fields is 283 kg/hectare (Qian et al., 2023), accounting for up to ∼11% (∼30 million metric tons) of total global CH_4_ emissions (Olivier and Peters, 2020), while N_2_O emissions from rice fields is 1.7kg/hectare account for 11% of global agricultural emissions (Islam et al., 2018; Win et al., 2020; Qian et al., 2023). CH_4_ sets the pace for warming in the near term as it traps very large quantities of heat over a shorter period. Hence, curbing CH_4_ emissions is one of the fastest and most effective strategies to reduce the rate of warming and limit temperature rise to 1.5°C. Several international organizations advocate strategies to reduce CH_4_ emissions from rice cultivation. Alternative agronomic practices have all been evaluated for their effectiveness in reducing CH_4_emissions (Yusuf et al., 2012; Bhatia et al., 2013; Xu et al., 2017; Oo et al., 2018; Liu et al., 2022; FAO 2023), however, the levels of reduction achieved are low, often affecting rice yield and crop robustness.

Here, we report data from a multi season open field study in rice with a methane-derived microbial biostimulant. There were three objectives with regards to the effect of methane-derived microbial biostimulant in paddy: (i) to assess the effect on grain yield improvement and reduction of CH_4_/N_2_O emissions; (ii) to understand the molecular mechanisms mediated by the microbial biostimulant in paddy; and (iii) to investigate the effect of reduced nitrogen (N) levels on grain yield and CH_4_/N_2_O flux. The study highlights a unique approach for achieving sustainable rice production and climate resiliency.

## 2. Materials and Methods

### 2.1 Field experimental design and cultivation practice

The field experiment to validate methane-derived microbial biostimulant was conducted at Vellore, Tamil Nadu, India, between June and October 2021 (season I) and February to June 2022 (season II). Field layout is shown in **Fig. S1a-b, supplementary materials.**

### 2.2 Microbial biostimulant application

The methane-derived microbial biostimulant (CleanRise™) is manufactured by String Bio, India, using an IP-protected fermentation process. The active ingredient in microbial biostimulant are cells of *Methylococcus capsulatus* derived by an innovative fermentation, downstream processing and formulation process (PCT application No. WO2021240472A1; Whole cell methanotroph based biostimulant compositions, methods and applications thereof). Two different treatment protocols were followed for season I study. With 100% NPK application, 10ml/L of microbial biostimulant was applied and with 75% N as input, three different doses of microbial biostimulant, 5ml/L (condition 1), 10ml/L (condition 2) and 15ml/L (condition 3) were tested. For season II, an optimal dose of microbial biostimulant at 10ml/L was evaluated under 100% NPK level. Elaborate experimental details are mentioned in **Supplementary methods** file. Grain yield in microbial biostimulant treated plots were compared with the respective control treatments and harvest index (HI) was computed following Du et al. (2022).

### 2.3 CH_4_ and N_2_O emission measurement

The static closed chamber method (Minamikawa et al., 2015) was used for gas sample collection in this study. For season I study, gas samples were collected at three time points [40, 60 and 80 days after transplanting (DAT) which respectively correspond to active tillering stage, panicle initiation stage and grain filling stage] while samples were collected every 10 days after transplantation during the season II evaluation. Gas samples were analyzed using gas chromatography with a Flame ionization detection (FID) and Thermal Conductivity Detector (TCD). CH_4_ and N_2_O flux were calculated and expressed as gram/hectare/hour (g/ha/h). The equivalent CO_2_ (CO_2_e) emission for total CH_4_ and N_2_O was calculated using the following Oo et al., 2018.

### 2.4 RNA extraction and transcript analysis

Total RNA extraction, cDNA synthesis and Reverse transcriptase-quantitative polymerase chain reaction (RT-qPCR) were carried out as described earlier (Kumar et al., 2018).

### 2.5 Statistical analysis

Average mean, standard error (SE) and number of replicates (n) used for each experiment were employed for statistical analysis using the GraphPad QUICKCALC online software (http://www.graphpad.com/quickcalcs/ttest1.cfm). The statistical significance of differences between controls and samples were tested according to the unpaired Student’s *t*-test.

Additional details about the methodology used in the study that are not detailed here are mentioned in the **Supplementary methods file**.

## 3. Results

### 3.1 Methane-derived microbial biostimulant improve growth and grain yield in paddy

To evaluate the effect of a methane-derived microbial biostimulant (CleanRise™) on rice grain yield, open field experiments were conducted across two seasons. With the application of microbial biostimulant, a significant increase in number of grains per spikelet and test weight was observed (**Table 1**). During the first season trial, the average grain yield improvement induced by microbial biostimulant varied between 32-39% (8004 ± 299 kg/ha to 8400 ± 80 kg/ha in microbial biostimulant treatment vs 6024 ± 216 kg/ha in control plots under 100% NPK levels) (**Fig. 1a** and **Fig. S2a, supplementary materials**). There was no significant difference between control and treatments with respect to straw yield (**Fig. 1b**). An informative indicator of the sink-source balance is the harvest index (HI). HI varies among rice varieties between 0.17-0.53 and further depends on environmental factors (Yang and Zhang, 2010). A HI of 0.39 was observed in response to microbial biostimulant application, while the HI observed for controls was 0.30 (**Fig. 1c**). During second season validation, microbial biostimulant application resulted in 39% improvement in grain yield (6997kg/ha in microbial biostimulant treatment vs 5015kg/ha in control plots) (**Fig. 1d**). Validation of microbial biostimulant in paddy in other testing locations also confirmed the positive impact of the methane-derived biostimulant across different seasons/ecological regions (**Fig. S2b-d, supplementary materials**).

**Figure 1.**
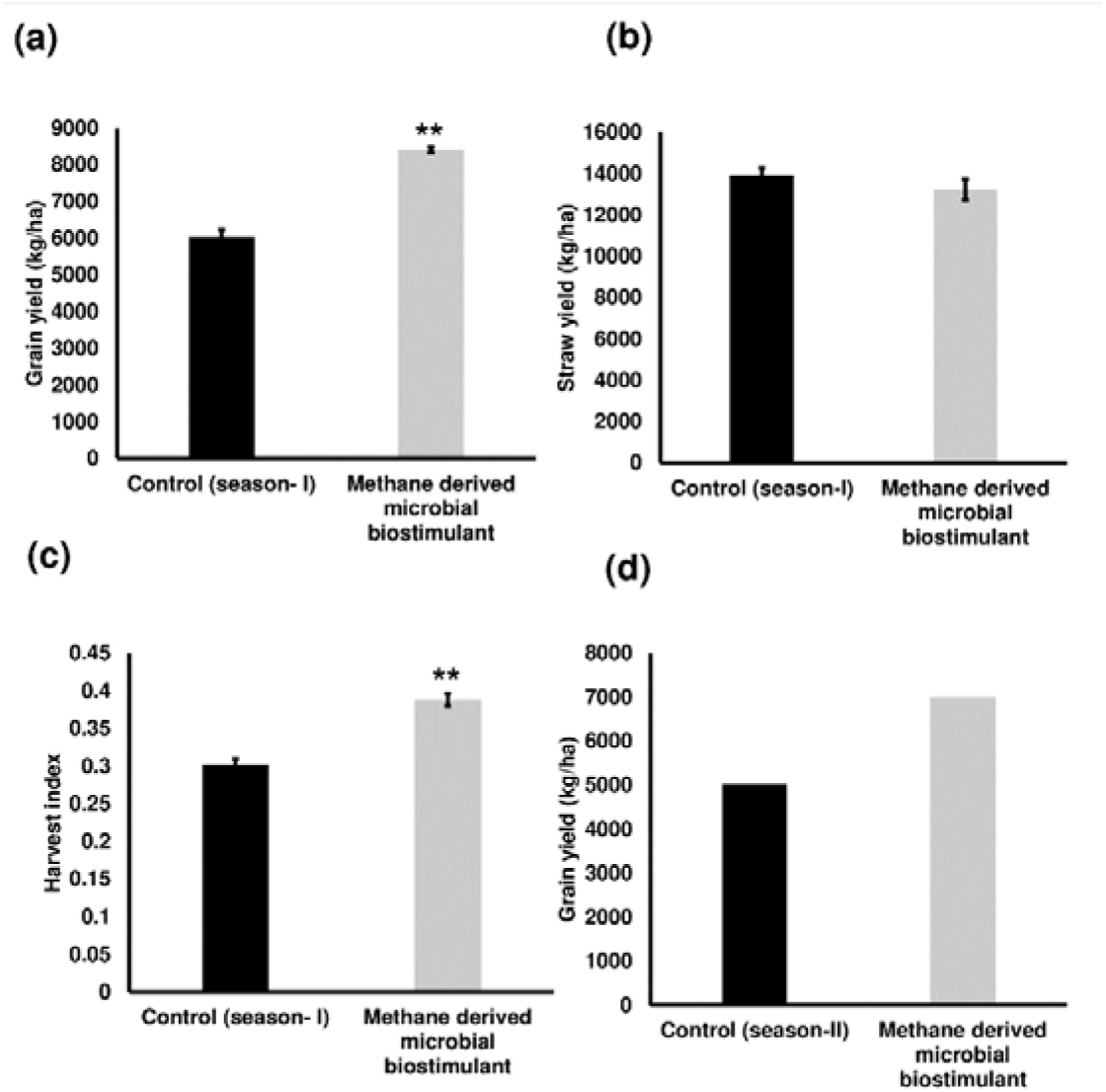
Methane-derived microbial biostimulant increases grain yield and harvest index in rice. (a) Effect of microbial biostimulant on grain yield improvement in paddy from season I validation-Methane derived microbial biostimulant application resulted in 39% improvement in grain yield compared to control. (b) Impact of methane derived microbial biostimulant on straw yield-There was no significant change in levels of straw yield between the treatments. (c) Impact of methane derived microbial biostimulant on harvest index in rice-A significant increase in harvest index of 0.39 was observed in microbial biostimulant treated plants compared to controls (0.30). (d) Influence of methane derived microbial biostimulant on grain improvement in paddy from season II validation-Grain yield improvement of∼ 39% was observed during second season validation. Control (season-I) and control (season-II) represents the yield observed in control plots from season I and season II validation respectively. As the second season trial was a demonstration trial in an area of ∼810 m2/treatment, bulk harvest was performed and hence error bar is not shown in the data. Differences were evaluated using the two-tailed Student’s *t* test and *P* < 0.01 is represented by “∗∗”.

**Table 1.** Grain yield component traits in methane derived microbial biostimulant treated paddy. Yield related traits mentioned are average data collected from 5 independent plants. Differences were evaluated using the two-tailed Student’s *t* test and *P* < 0.05 and *P* < 0.01, and *P* < 0.001 are represented by “∗” and “∗∗”.

| Treatment | Grains/spikelets | Test weight (g) | Grain yield (kg/ha) | Yield improvement (%) |
| --- | --- | --- | --- | --- |
| **Control**<br>**(season-I)**<br>**100% N** | 128±6.7 | 22±0.86 | 6024±216 | 0 |
| **Microbial**<br>**Biostimulant**<br>**100% N** | 166±7.6<br>* | 28±1.12<br>* | 8400±80<br>** | 39.44 |

### 3.2 Microbial biostimulant regulate photosynthesis, tillering and panicle architecture in paddy

Phenotypic analysis in microbial biostimulant treated paddy leaves showed brilliant dark-green leaves compared to control leaves (**Fig. S3a, supplementary materials**). We observed that microbial biostimulant application resulted 18% increase in photosynthetic rate, 22% increase in stomatal conductance and ∼48% increase in transpiration rate (**Fig. S3b-d, supplementary materials**). To elucidate the molecular mechanism affecting the phenotype, we carried out mRNA expression analysis of genes encoding enzymes involved in photosynthesis, tillering and panicle architecture. The transcript levels in microbial biostimulant treated leaves or panicles were compared with respect to control samples. Most of the genes related to photosynthesis were upregulated between 1.4-fold to ∼20-fold in plants applied with microbial biostimulant. The up-regulated genes were related to all major components of photosynthesis, including, chlorophyll biosynthesis pathway and chloroplast development, Photosystem I, Photosystem II and enzymes involved in the CBB cycle (Calvin-Benson-Bassham) (**Fig. 2a-b**). This data suggests that microbial biostimulant application positively influences photosynthesis through up-regulation of specific targeted pathways (**Fig. 2a-b**).

**Figure 2.**
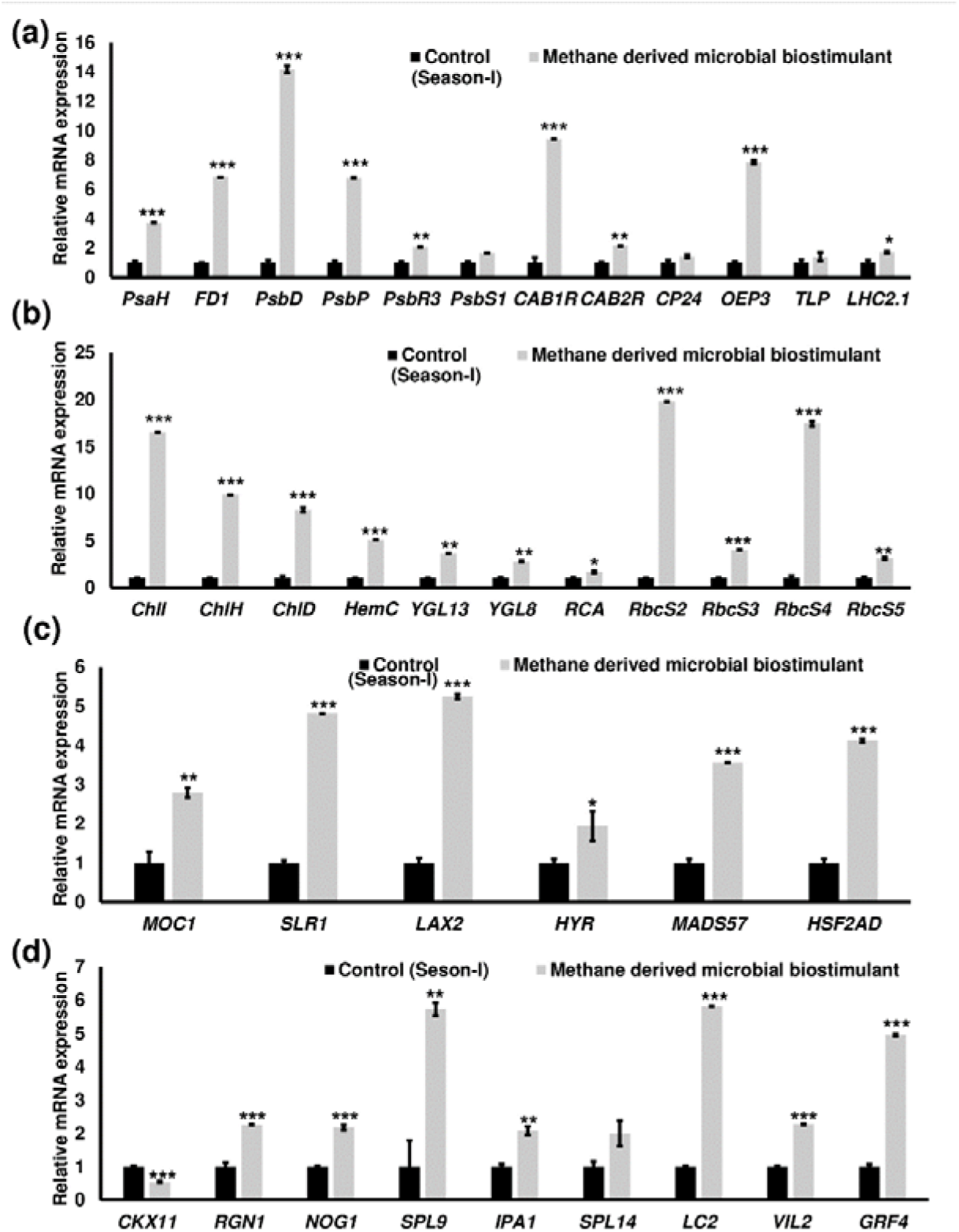
Microbial biostimulant acts as a master regulator of photosynthesis, tillering and panicle architecture. Reverse transcriptase-quantitative polymerase chain reaction (RT-qPCR) analysis showing the relative expression of in genes related to photosynthesis (a & b), tillering (c) and panicle architecture (d) in rice with or without microbial biostimulant application. Expression levels of genes were normalized to the endogenous reference gene *actin* and are represented relative to respective controls, which was set to 1. Pooled leaves or panicles from three to five plants were used for RNA extraction. The results shown are from three independent experiments. **Abbreviations:** Photosystem I reaction center subunit VI (*PsaH*); Ferredoxin 1(*FD1*); Photosystem II D2 protein (*PsbD*); photosystem II subunit P (*PsbP*); photosystem II subunit PsbR3 (*PsbR3*); Photosystem II 22 kDa protein 1 (*PsbS1*); Chlorophyll a-b binding protein 1 (*CAB1R*); Chlorophyll a-b binding protein 2 (*CAB2R*); Chlorophyll Protein 24 (*CP24*); Oxygen-evolving enhancer protein-3 (*OEP3*); Thylakoid luminal protein (*TLP*); Chlorophyll a-b binding protein 2.1/Light Harvesting Complex Protein 2.1 (*LHC2.1*); Magnesium-chelatase subunit ChlI (*ChlI*); Magnesium-chelatase subunit ChlH (*ChlH*); Magnesium-chelatase subunit Child (*<u>ChlD</u>*); porphobilinogen deaminase/ hydroxymethylbilane synthase (*HemC*); yellow-green leaf 13 (*YGL13*); yellow-green leaf 8 (*YGL8*); Rubisco activase (*RCA*); Ribulose bisphosphate carboxylase small subunit (*RbcS2,3,4,5*); monoculm 1 (*MOC1*); Slender Rice-1 (*SLR-1*); LAX PANICLE2 (*LAX2*); HIGHER YIELD RICE (*HYR*); MADS-box transcription factor (*MADS57*); Heat Stress Transcription Factor 2D (*HSF2AD*); Cytokinin oxidase/dehydrogenase (*CKX11*); Regulator of Grain Number-1 (*RGN1*); Number of Grains-1 (*NOG1*), SQUAMOSA Promoter Binding Protein-Like (*SPL9* and *SPL14*), Ideal Plant Architecture-1 (*IPA1*); Leaf Inclination 2/VIN3 (vernalization insensitive 3-like protein)-(*LC2*); VIN3-LIKE 2 (*VIL2*); Growth Regulating Factor 4 (*GRF4*); Differences were evaluated using the two-tailed Student’s *t* test and significant differences at *P* < 0.05, *P* < 0.01, and *P* < 0.001 are represented by “∗” “∗∗”, and “∗∗∗”, respectively.

To further understand the molecular mechanism of enhanced photosynthetic capacity on axillary meristem growth and panicle architecture, next we examined the transcript levels of critical genes involved in regulation of shoot branching, panicle and grain development. There was an enhanced expression of tillering related genes ranging from 1.9-fold to 5.2-fold by microbial biostimulant application (**Fig. 2c**). As photosynthate partitioning from source (leaf) to the sink (grains) is critical for panicle development and grain filling, mRNA expression of key genes involved in grain development were further analyzed. A 2-fold to 5.7-fold upregulation of genes controlling panicle architecture was observed indicating that improved photosynthetic capacity positively translated to grain filling and development (**Fig. 2d**). Interestingly, microbial biostimulant application also downregulated the expression of *CKX11*, a negative regulator of panicle architecture in paddy. The above results provide evidence that microbial biostimulant acts as a major regulator of multiple systemic pathways that improve photosynthesis, higher number of productive tillers and better panicles thus resulting in superior yield.

To assess the impact of microbial biostimulant on nutrient uptake, we investigated the effect on expression of key genes encoding macronutrient transporters. Our reverse transcription-quantitative polymerase chain reaction (RT-qPCR) analyses showed the upregulation of genes involved in nitrogen uptake and transport by 2-fold to 12-fold in microbial biostimulant treated paddy roots (**Fig. S4a, supplementary materials)**. Further, gene expression analysis of high affinity potassium and phosphate transporters also indicated a 2-fold to 10-fold increase in microbial biostimulant treated roots (**Fig. S4b, supplementary materials**). The data indicates a direct influence of the microbial biostimulant on nutrient uptake and utilization, particularly nitrogen, phosphate and potassium.

### 3.3 GHG mitigation potential of methane-derived microbial biostimulant

We next studied the effect of microbial biostimulant on flux of CH_4_ and N_2_O from rice paddies during three time points of crop growth (40, 60, 80 DAT) of season I study. The dynamic fluxes of CH_4_ and N_2_O over the rice growing period were strongly affected by the microbial biostimulant application. In our studies, CH_4_ and N_2_O flux were high during the tillering stage, then gradually decreased towards the flowering stage and end of the growing period across all the plots (**Fig. 3**). CH_4_ emission varied considerably among the treatments and the dynamics of CH_4_ flux during the cropping seasons is presented in **Fig. 3a**. Microbial biostimulant application resulted in a reduction of approximately 70% in CH_4_ emissions at 40 DAT (46± 3.78 g/ha/h CH_4_ in microbial biostimulant treated plants vs 176± 9.65 g/ha/h CH_4_ in control plants). Approximately 50% reduction in emission was recorded during subsequent sampling at 60 DAT (29.3± 1.58 g/ha/h CH_4_ in microbial biostimulant treated plants vs 59.2 ± 1.3 g/ha/h in control plants) and 80 DAT (14± 1.30 g/ha/h CH_4_ in microbial biostimulant treated plants vs 31.36 ± 0.31 g/ha/h in control plants). Although the levels of N_2_O emissions were much lower compared to CH4 flux, a similar emission pattern was observed. Fluxes of N_2_O at the farms varied from 2.3 g/ha/h to 5.7 g/ha/h in microbial biostimulant treatment, compared to 4.2 g/ha/h to 8.2 g/ha/h in control plots (**Fig. 3b**). Highest N_2_O flux was 8.26± 0.23 g/ha/h during early crop growth in control plants. Here, microbial biostimulant application led to a significant reduction in N_2_O emission upto 30% (5.7<u>6+</u> 0.29 g/ha/h). Microbial biostimulant-mediated reduction in N_2_O flux was in the range of ∼45% during the second and third sampling periods. Cumulative CH_4_ and N_2_O emissions from the rice field during the overall rice-growing season showed significant differences between all treatments. Average CH_4_ emission during first season cropping recorded from the control plots was 244.10 kg/ha and microbial biostimulant application reduced it to 80.2043kg/ha, leading to 67% reduction in CH_4_ flux (**Table 2**). Similarly, there was ∼35% cumulative reduction in N_2_O flux with microbial biostimulant application, when compared with N_2_O flux from the control plot (**Table 2**). During phase II trials, although there was no significant change in CH_4_ emission levels at 10 and 20 DAT, there was a peak reduction ranging from 23%-50% during the subsequent sampling period (**Fig. 3c**). N_2_O reduction during season II varied between 30-70% during the crop growth (**Fig. 3d**). Taken together, from two season trials, this study provides conclusive evidence that significant improvement in grain yield with concomitant reduction in GHG emission from paddy cultivation can be enabled with methane derived microbial biostimulant.

**Figure 3.**
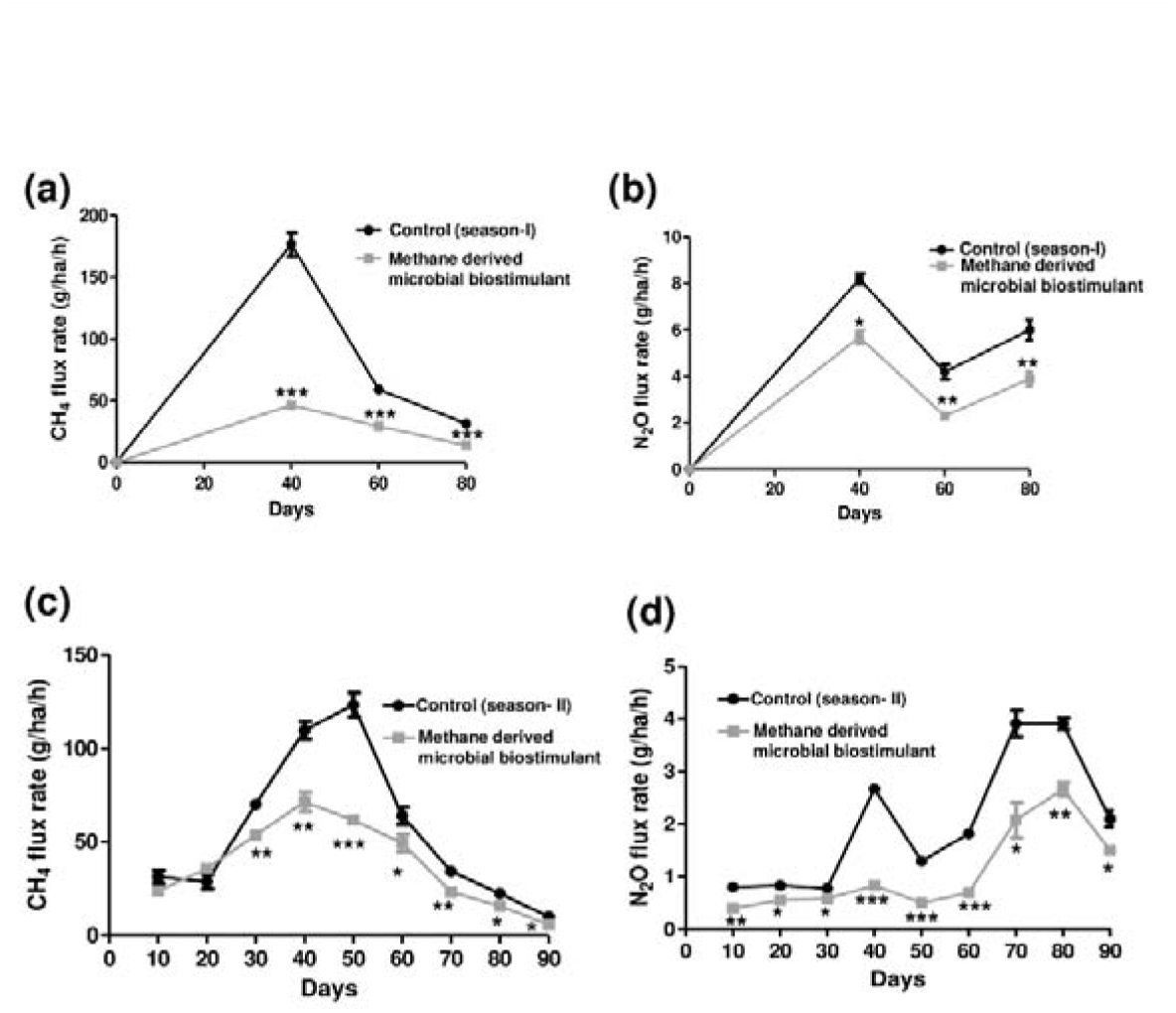
Greenhouse gas mitigation mediated by methane-derived microbial biostimulant-Influence of microbial biostimulant on CH_4_ and N_2_O emission from rice field-Gas samples were collected in triplicate from each plot for every time point and were analyzed using gas chromatography with a thermal conductivity detector (TCD). While gas samples were collected at three time points for season I trials, gases were collected at every 10 days during season II trial. CH_4_ and N_2_O flux were calculated and expressed as gram/hectare/hour (g/ha/h). Microbial biostimulant application resulted in ∼50- >70% reduction in CH_4_ (a) emission from rice fields whereas it was between 30-45% reduction in N_2_O (b) during the course of plant growth during season I study. Methane-derived microbial biostimulant use resulted in∼35% reduction in CH_4_ (c) emission from rice fields whereas it was between ∼50% reduction in N_2_O (d) during season II validation. Control (season-I) and control (season-II) represent the emission observed in control plots from season I and season II validation respectively. Differences were evaluated using the two-tailed Student’s *t* test and *P* < 0.05, *P* < 0.01, and *P* < 0.001 are represented by “∗”, “∗∗”, and “∗∗∗”, respectively.

**Table 2.** Effect of methane-derived microbial biostimulant on yield-scaled CO2-eq emission in rice. Yield-scaled CO2e-emission of CH4 and N2O were found to be significantly lower in methane-derived microbial biostimulant treatment compared to controls.

| Treatments | Methane emission |  |  | Nitrous oxide emission |  |  |
| --- | --- | --- | --- | --- | --- | --- |
|  | Cumulative emission (kg/ha) | CO<sub>2</sub>-eq emissions (kg CO<sub>2</sub>/ha) | Yield-scaled CO<sub>2</sub>-eq emission (kg CO<sub>2</sub>-eq/t) | Cumulative emission (kg/ha) | CO<sub>2</sub>-eq emissions (kg CO<sub>2</sub>/ha) | Yield-scaled CO<sub>2</sub>-eq emission (kg CO<sub>2</sub>-eq/t) |
| Control (100% N) | 244.10 | 20504.40 | 3403.78 | 18.50 | 5513.00 | 861.40 |
| Microbial Biostimulant | 80.2 0 | 10432.80 | 802.00 | 11.90 | 3546.20 | 422.16 |
| Control<br>(75% N) | 235.00 | 19740.00 | 3589.09 | 13.84 | 4123.33 | 749.70 |
| Microbial<br>Biostimulant<br>condition- 1 | 130.71 | 10980.20 | 1542.16 | 10.34 | 3081.32 | 432.77 |
| Microbial<br>Biostimulant<br>condition- 2 | 87.86 | 7380.80 | 997.41 | 6.76 | 2015.47 | 270.90 |
| Microbial<br>Biostimulant<br>condition- 3 | 44.97 | 3777.76 | 490.62 | 6.23 | 1855.46 | 240.97 |

### 3.4 Microbial biostimulant improved paddy grain yield and lowered GHG emission under reduced nitrogen inputs

The combination of high-yielding crop varieties and the widespread use of inorganic fertilizers has markedly improved crop production. However, excessive Nitrogen (N) input can lead to severe environmental pollution. As optimizing nitrogen management is among the promising avenues to reduce GHG emissions from rice paddies and the fact that microbial biostimulant application modulated genes of N uptake and transport (**Fig. S4a, supplementary materials**), we tested the effect of microbial biostimulant on yield and GHG emission by reducing the N fertilizer level. A 25% reduction in N fertilizer levels decreased grain yield in the control treatment (75% N control), whereas reduced N application combined with microbial biostimulant treatment improved grain yield significantly without altering straw yield (**Fig. 4a-b**). On average, grain yields under reduced N, ranged between 28%-38% with different doses of microbial biostimulant. The maximum and minimum rice grain yield under reduced N was 7714±399.14 kg/ha with microbial biostimulant condition-3 and 7129 ± 589.63 kg/ha with microbial biostimulant condition-1 respectively compared to 5561 ± 253.24 kg/ha for control (75%N) plants. It has been reported that N applications at basal and tillering stages are important for improved tillering and increased panicle number to ensure high yield (Kamiji et al., 2011). Microbial biostimulant application could partially reduce the need for the exogenous supply of N by improving the plants’ NUE.. Moreover, reduced N application resulted in improved root growth in microbial biostimulant treated plants compared to control plants (**Fig. S5a, supplementary materials**). Microbial strains in biostimulant produced 1.83-3.61mg/L indole acetic acid (IAA) in presence of tryptophan (**Table S1, supplementary materials**). With reduced N, the HI was also significantly enhanced by microbial biostimulant application and among the three conditions tested, the maximum HI of 0.39 was recorded with condition-2 (**Fig. S6, supplementary materials**).

**Figure 4.**
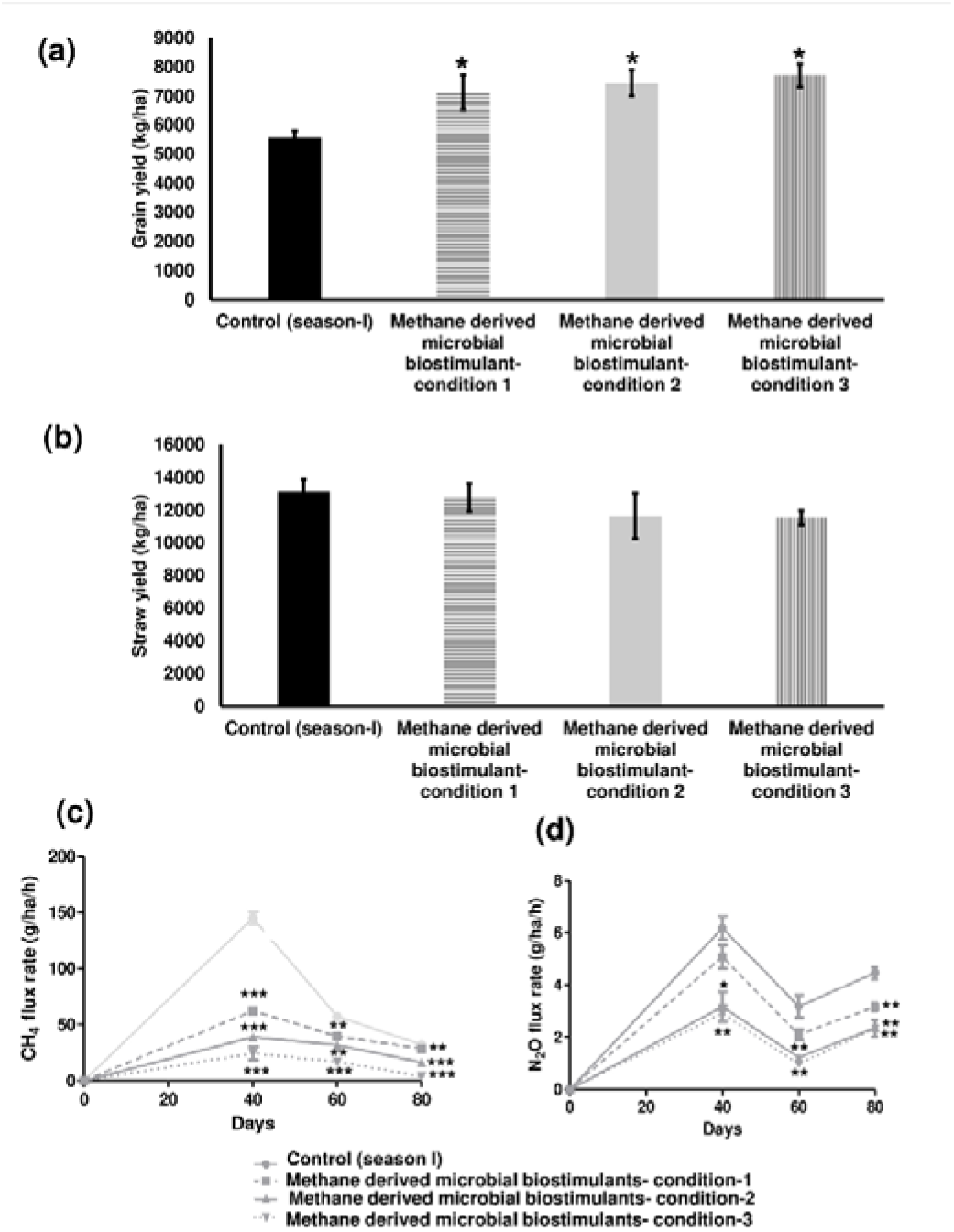
Microbial biostimulant mediated grain yield improvement and greenhouse gas (GHG) emission reduction in rice under reduced nitrogen (N) level. (a) Effect of microbial biostimulant on grain yield improvement in paddy with 75% N fertilizer-Methane-derived microbial biostimulant application resulted in 28-39% improvement in grain yield with different doses (condition 1-5ml/L, condition 2-10ml/L and condition 3-15ml/L) of microbial biostimulants compared to 75% N control. (b) Impact of methane derived microbial biostimulant on straw yield-There was no significant change in levels of straw yield between the treatments. (c) and (d) Impact of microbial on CH_4_ and N_2_O emission from rice-Gas samples were collected in triplicate during three critical phase of plant growth and were analyzed using gas chromatography with a thermal conductivity detector (TCD). CH_4_ and N_2_O flux were calculated and expressed as gram/hectare/hour (g/ha/h). Methane derived microbial biostimulant application resulted in 45->70% reduction in CH_4_ flux and 26->40% reduction N2O emission during the course of rice growth. g/ha/h-gram/hectare/hour. Significant differences at *P* < 0.05, *P* < 0.01, and *P* < 0.001 are represented by “∗”, “∗∗”, and “∗∗∗”, respectively.

A dose dependent change in CH_4_ and N_2_O flux was observed with different microbial biostimulant conditions under reduced N levels. Maximum CH_4_ flux of 145± 5.64 g/ha/h was recorded in the controls whereas the lowest flux of 24.5± 6.02 g/ha/h was observed in microbial biostimulant condition-3 at 40 DAT (**Fig. 4c**). CH_4_ flux with microbial biostimulant condition-1 and condition-2 was observed at 62.21± 1.69 g/ha/h and 39± 2.64 g/ha/h respectively. Thereafter, the flux decreased to 57.3<u>6+</u> 1.41 g/ha/h in control plants and 39.8 ± 1.66 g/ha/h, 31.9 ± 2.81 g/ha/h and 16.50 ± 2.33 g/ha/h in microbial biostimulant treated plants during the second sampling stage. As maturity stage approached, the CH_4_ flux further declined to 32.46± 0.69 g/ha/h in control plants and ranged between 28.7± 0.37 g/ha/h, 16.96 ± 1.03g/ha/h and 3.96± 0.92 g/ha/h with microbial biostimulant application. Overall, the CH_4_ flux was considerably lower in microbial biostimulant treated plots than in control plots. The N_2_O flux under reduced N application varied between 6.19g/ha/h-3.18g/ha/h in control (**Fig. 4d**). With different treatments of the microbial biostimulant, lower N_2_O flux was observed under all the three conditions. At 40 DAT, N_2_O flux from control was 6.19+ 0.44 g/ha/h while microbial biostimulant treatment resulted in 5.07± 0.44 g/ha/h, 3.16± 0.56g/ha/h and 2.91± 0.30 g/ha/h. Subsequently, at 60DAT control plants recorded 3.18± 0.42 g/ha/h and microbial biostimulant treatment showed N_2_O flux ranging from 2.10 ± 0.20 g/ha/ha, 1.2<u>2+</u> 0.09g/ha/h and 0.97± 0.09 g/ha/h. Towards the maturity stages, N_2_O flux from control was 4.46± 0.23 g/ha/h and microbial biostimulant treatment recorded 3.16± 0.16 g/ha/h, 2.37± 0.13g/ha/h and 2.34<u>+0.32g/ha/h</u>. Overall, the results clearly indicated that cumulative CH_4_ and N_2_O emissions were significantly lower when N input reduction was combined with microbial biostimulant application (**Table 2**). This study demonstrates a powerful way to reduce GHG emissions from rice fields while permitting savings on fertilizers and increased crop yields by the application of a methane-derived microbial biostimulant.

### 3.5 Impact on yield-scaled CO_2_ reduction mediated by methane-derived microbial biostimulant

The impact of GHG emission is quantitatively assessed by computing global warming potential (GWP) that accounts for all sources (carbon and non-carbon) of CO_2_e (Robertson et al., 2000; Mosier et al., 2006). In the present study, the contribution of CH_4_ to the total GWP ranged from ∼3777 kg CO2/ha to 20504 kg CO2/ha under the different treatments. Yield-scaled CO_2_ equivalent of CH_4_ emission from controls were 3403 kg CO2-eq/t whereas there was significant reduction of 802 kg CO2-eq/t with microbial biostimulant application under 100% N fertilizer level **(Table 2)**. Similarly, N_2_O equivalent CO_2_ emission from fields with microbial biostimulant application was only 422 kg CO2-eq/t compared to 861 kg CO2-eq/t from control fields **(Table 2)**. High level of N2O emission could be possibly due to gas sampling after 3-4 days of top dressing with N fertilizer. Reduced N inputs combined with microbial biostimulant application also impacted CO_2_eq emissions. Difference in the CH_4_ equivalent CO_2_e emission under reduced N inputs ranged from 1542-490-kg CO2-eq/t among different conditions of microbial biostimulant tested compared to ∼3589 kg CO2-eq/t in controls. While N_2_O equivalent CO_2_ emission was as high as 750 kg CO2-eq/t in controls the levels varied between ∼240-432 kg CO2-eq/t with different microbial biostimulant conditions. Overall, microbial biostimulant application reduced the yield-scaled GWP by upto 77% and 50% of CH_4_e and N_2_Oe share respectively over control fields. Interestingly, microbial biostimulant application along with reduced N inputs recorded significant reduction in CO2e emission between ∼60% to >80% for CH_4_eq and 41%-60% for N_2_Oeq (**Table 2**).

### 4.0 Discussion

The demand for increased agricultural production in the context of arable land scarcity and climate change needs innovative solutions to overcome challenges and address inefficiencies. Global rice consumption has increased markedly, growing from157 million tonnes in 1960 to 520 million tonnes in 2022 (USDA, 2023). Global rice demand is further projected to increase by 28% in 2050, yet rice yields have stagnated in 35% of all rice-growing regions (Ray et al., 2012). Here, through multi-location and multi-season trials, we demonstrate a substantial increase in grain yield ranging from 15-39% (**Fig. 1a and Fig. S2, supplementary materials**) with methane-derived microbial biostimulant. Methane-derived microbial biostimulant (CleanRise™) is a promising solution to enhance the yield potential in rice to satisfactorily address global food security. A significant additional effect of microbial biostimulant application is the improved NUE observed under reduced N application levels (**Fig. 3a and Fig S7a-b, supplementary materials**). Even with 25% reduced N application, the yield per hectare was enhanced over the control (75%N) treatment. While the optimal requirement of N may vary with soil condition and crop management, the study demonstrates that a similar approach could be considered for exhaustive cereal crops like maize and wheat.

It is interesting to note that methane-derived microbial biostimulant application resulted in significantly improved root growth and enhanced photosynthetic capacity per unit leaf area, which further translated into higher panicle number and test weight, and thus superior rice yield (**Table 1, Figs. S3 & S8, supplementary materials and Fig. 1a & 1d**). We established that *M. capsulatus* in the microbial biostimulant formulation were able to symbiotically associate with root and leave tissues of paddy (**Fig. S9**) and have a significant effect on host transcriptional regulation (**Fig. 5**). Based on the phenotypic and genotypic observations, we propose three major routes for mode of action of microbial biostimulant in rice. First, microbial biostimulant positively regulated multiple pathways related to macronutrient availability, uptake and transport, resulting in better nutrient use efficiency (**Fig. S4a-b, Figs. 5 and 7**). Secondly, microbial cells were able to produce and hence supply IAA to plants thus accelerating auxin mediated root growth and crop establishment (**Table S1**). Third, microbial biostimulant simultaneously regulated diverse pathways regulating photosynthesis, axillary branching and panicle development (**Figs. 2 and 5 & Table 1**). Often, there are several check points to regulate photosynthesis and carbon partitioning in plants (Paul and Foyer, 2001). We propose that microbial biostimulant mediated enhanced gene expression along with superior photosynthetic activity translated to improved axillary bud initiation and carbon fixation. Further, effective photosynthate partitioning to sink tissues, like flag leaves, panicles and developing grains, possibly translated to better yield. Ambavaram et al. (2014) also reported efficient translocation of carbohydrates from source to sink to improve grain yield in paddy. It has been previously reported that even a minor increase in net photosynthetic activity translated to better yield in wheat and rice (Parry et al., 2011; Li et al., 2020). It is interesting to note that the methane-derived microbial biostimulant enhanced the expression of positive regulators and downregulated negative regulator in paddy to improve crop performance and yield (**Figs. 2d and 5**). Our finding systematically highlights the in-depth molecular mechanisms mediated by biostimulant with modulation of critical physiological events like photosynthesis, tillering and panicle formation in rice.

**Figure 5.**
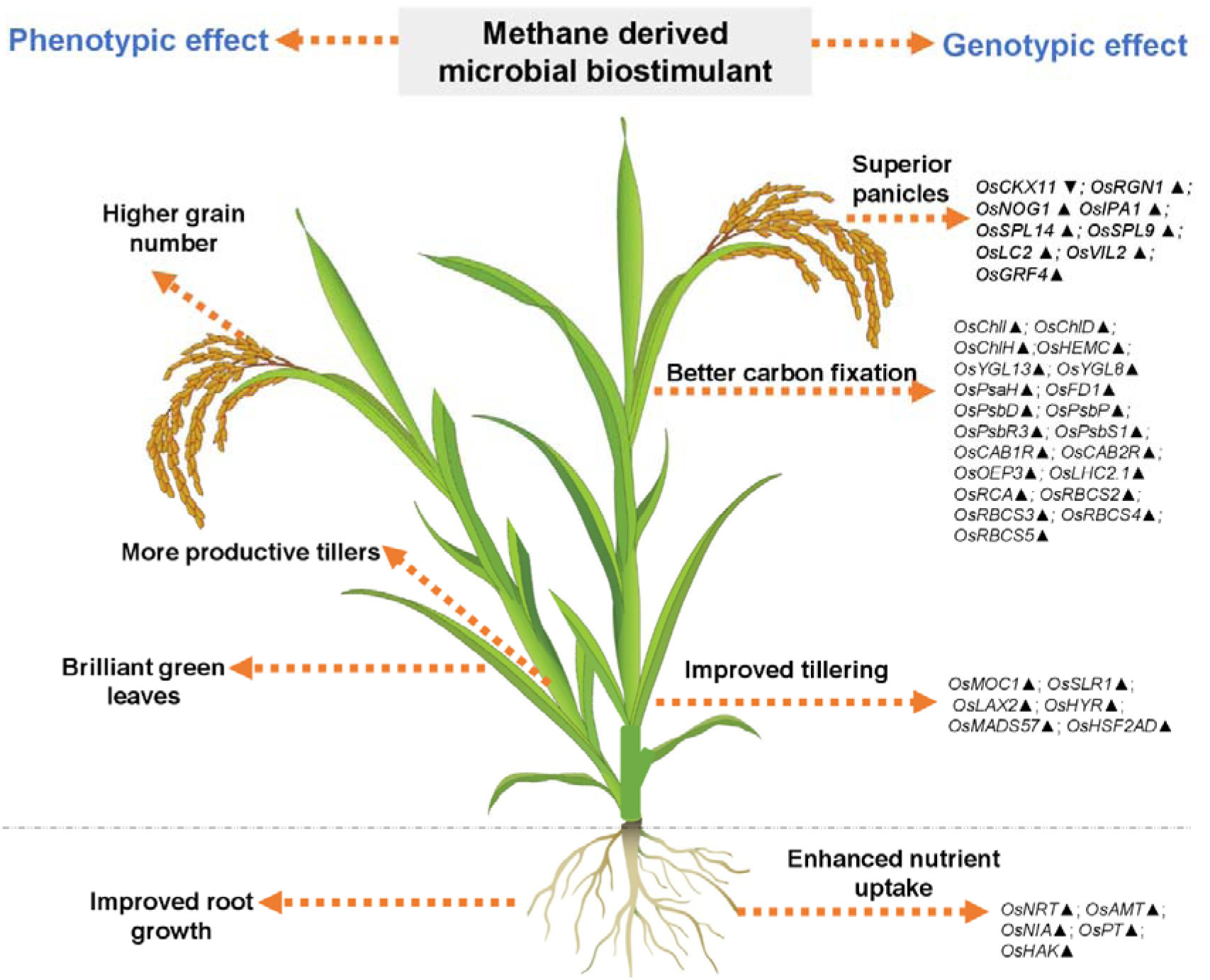
Representative image showing overview of phenotypic and genotypic traits modulated by microbial biostimulant application in rice (*Oryza sativa*). Microbial cells in the biostimulant formulation improve macronutrient availability and transport. Microbial biostimulant application modulates expression of gene involved in axillary bud formation resulting in more productive tillers. Targeted activation of genes related to chlorophyll biosynthesis pathway and chloroplast development, Photosystem I, Photosystem II and CBB cycle (Calvin-Benson-Bassham) results in improved carbon fixation. Active photosynthate translocation to developing grain and biostimulant mediated activation of genes involved in panicle architecture results in a greater number of grains per panicle translating to superior yield. **Abbreviations**: *Oryza sativa (Os);* Nitrate transporter (NRT); Ammonium transporter (AMT); Nitrate reductase (NIA); Phosphate transporter (PT); High affinity potassium transporter (KKT); Photosystem I reaction center subunit VI (*PsaH*); Ferredoxin 1(*FD1*); Photosystem II D2 protein (*PsbD*); photosystem II subunit P (*PsbP*); photosystem II subunit PsbR3 (*PsbR3*); Photosystem II 22 kDa protein 1 (*PsbS1*); Chlorophyll a-b binding protein 1 (*CAB1R*); Chlorophyll a-b binding protein 2 (*CAB2R*); Chlorophyll Protein 24 (*CP24*); Oxygen-evolving enhancer protein-3 (*OEP3*); Thylakoid luminal protein (*TLP*); Chlorophyll a-b binding protein 2.1/Light Harvesting Complex Protein 2.1 (*LHC2.1*); Magnesium-chelatase subunit ChlI (*ChlI*); Magnesium-chelatase subunit ChlH (*ChlH*); Magnesium-chelatase subunit Child (*<u>ChlD</u>*); porphobilinogen deaminase/ hydroxymethylbilane synthase (*HemC*); yellow-green leaf 13 (*YGL13*); yellow-green leaf 8 (*YGL8*); Rubisco activase (*RCA*); Ribulose bisphosphate carboxylase small subunit (*RbcS*); monoculm 1 (*MOC1*); Slender Rice-1 (*SLR-1*); LAX PANICLE2 (*LAX2*); HIGHER YIELD RICE (*HYR*); MADS-box transcription factor (*MADS57*); Heat Stress Transcription Factor 2D (*HSF2AD*); Cytokinin oxidase/dehydrogenase (*CKX11*); Regulator of Grain Number-1 (*RGN1*); Number of Grains-1 (*NOG1*), SQUAMOSA Promoter Binding Protein-Like (*SPL9* and *SPL14*), Ideal Plant Architecture-1 (*IPA1*); Leaf Inclination 2/VIN3 (vernalization insensitive 3-like protein)-(*LC2*); VIN3-LIKE 2 (*VIL2*); Growth Regulating Factor 4 (*GRF4*). Upward arrow (▴) indicates gene upregulation more than 1.5 fold and downward arrow (▾) indicates more than 50% downregulation of genes.

The field experiments also clearly demonstrate significantly reduced CH_4_ and N_2_O emissions, both with standard and reduced N levels with microbial biostimulant treatment (**Figs. 3 and 4**). Primarily, rice plants serve as the major conduits for the transfer of CH_4_ from the soil to the atmosphere. A well-developed aerenchyma cells in leaf blade, sheath, culm and roots of rice plants makes a good passage for the gas exchange between the atmosphere and the soil (Nouchi et al., 1990; Nouchi and Mariko 1993; Friedl et al., 2010; Li et al., 2013). A majority of CH_4_ (∼90%) formed in rice soil and is emitted through aerenchyma in rice plants by the process of diffusion (Bhattacharyya et al., 2019). Also, rice paddy utilizes one-seventh of N fertilizer, making a more potent zone of N_2_O formation and emissions. With the observed symbiotic association in plants (**Fig. S9, supplementary materials**), it is highly plausible that the methane-derived microbial biostimulant carries out methane oxidation and thus utilize the methane for their growth. Similarly, the observed reduction in N_2_O emission from rice could be attributed to improved NUE mediated by microbial biostimulant (**Fig. S7, supplementary materials**).

Although rice is the main staple food for nearly half the world’s population, rice cultivation contributes to 283 kg/ha and 1.7 kg/ha respective to CH_4_ and N_2_O emissions annually (Qian et al., 2023). Rice growing economies are also among the leading methane emitters globally. For instance, countries like China, India and Indonesia have the largest rice cultivation area and contribute to 22-38%, 11-19% and 7-9% of the 24–37ℒTg per year global total, respectively (FAO, 2022; EDGAR v7.0. Global Greenhouse Gas Emissions, 2022). To meet the net zero targets, an ideal goal for different nations now is to reduce short- and long-term emissions without compromising crop yield. Currently, only 1/5^th^ of countries (25/148) mention rice mitigation measures in nationally determined contributions to the Paris Agreement (Rose et al., 2021). Here, we provide science-based solutions to prioritize actions to reduce agricultural CH_4_ emissions. At the COP26 meeting, countries aligned to a 2% reduction target in CH_4_ annually and the data outlined here highlights a powerful path to help achieve these targets. For instance, microbial biostimulant application to just 10% of the global paddy-cultivation area (16.2 million hectares) could deliver up to 24% of the global CH_4_ reduction target. Use in 30% of paddy cultivation area (48.6 million hectare) could help to achieve 72% of the global CH_4_ emission target. More ambitiously, enabling use in 50% of the worlds’ paddy cultivated area (81 million hectares) could deliver 120% of the reduction target (**Graphical abstract & Table S2**). Use of single disruptive solution like methane derived microbial biostimulant (CleanRise) thus could form a promising strategy to curb global CH_4_ emissions from farmed rice while meeting the COP26 target.

## 5. Conclusion

Reconciling rapidly increasing food demand with the need to address climate change by reducing emissions from agriculture is a complex problem requiring novel policy measures to incentivize best practices. Our study shows that use of methane derived microbial biostimulant is a win-win solution to improve yield, optimize NUE and reduce GHG emissions from rice fields. It provides the means to achieve the intensification necessary to address the food security for a growing world population, without compromising environmental and climate mitigation strategies. The mitigation pathways and optionality highlighted in this study can be accelerated with targeted policies and catalyze sustainable rice cultivation across the globe to address food security.

## Supplementary Figures

**Supplementary Fig 1a-** Experimental field layout for season I testing-Treatment details are below: T1- Control (100% NPK); T3 & T4- 10 ml/L dose of methane derived microbial biostimulant (100% NPK); T5- Control (75%N); T6- 75%N+microbial biostimulant 5ml/L (condition 1); T7- 75%N+ microbial biostimulant 10ml/L (condition 2); T8- 75%N+ microbial biostimulant 15ml/L (condition 3); T2, T9 and T10 are outside purview of this manuscript and hence are not discussed/explained. R1, R2 and R3 respectively corresponds to replication 1, 2 and 3. Small green box indicate position of gas collection base & chambers.

**Supplementary Fig 1b- Experimental field layout for season II testing** - Treatment details are below: T1- Control (100% NPK); T2- Microbial biostimulant- 10ml/L (100%NPK). Small square box indicate position of gas collection base & chambers.

**Supplementary Figure 2a-** Influence of methane derived microbial biostimulant on grain yield where second application was given as foliar spray instead of soil spray.

**Supplementary Figure 2b-d- Multilocation microbial biostimulant validation data**- Grain yield improvement mediated by methane derived microbial biostimulant under different agro ecological locations in India. Differences were evaluated using the two-tailed Student’s *t* test and significant differences at *P* < 0.05 and *P* < 0.01 are represented by * and ** respectively.

**Supplementary Fig 3(a)- Phenotypic feature of microbial biostimulant treated paddy leaves-** Influence of methane derived microbial biostimulant on greenness in paddy leaf : Control leaf (C) and methane derived microbial biostimulant treated leaf (MB).

**Supplementary 3(b-d)- Effect of methane derived microbial biostimulant on physiological traits in paddy leaves**- Photosynthetic efficiency, stomatal conductance and transpiration rate are represented as % relative to control plants. Student’s *t-*test: significant differences at *P* < 0.05 and *P* < 0.01 are represented by * and ** respectively

**Supplementary Fig 4- Influence of methane derived microbial biostimulant on expression of root nutrient uptake and transporter genes.**

RT-qPCR analysis showing the expression of genes related to macronutrinet transport and metabolism in roots of microbial biostimulant treated plants. Expression levels of genes were normalized to the endogenous reference gene actin and are represented relative to respective control roots, which was set to 1. Pooled root samples from control and microbial biostimulant treated roots used for RNA extraction. The results shown are from three independent experiments. Error bars indicate mean ± SE. Student’s t-test: significant differences at *P* < 0.05, *P* < 0.01 and *P* < 0.001 are represented by *, ** and ***, respectively. Nitrate transporter (NRT); Ammonium transporter (AMT); nitrate reductase (NIA); Glutamine synthetase (GS); glutamate synthase (GOGAT); Phosphate transporter (PT); High affinity potassium transporter (HAK); Zinc transporter (ZIP).

**Supplementary Fig 5- Effect of microbial biostimulant on root length-** Seedling root dipping was performed in paddy roots with microbial biostimulant and twenty days after transplanting seedlings were uprooted and root length was measured. Student’s t-test: significant differences at *P* < 0.01 is represented by “**”.

**Supplementary Fig 6- Impact of methane derived microbial biostimulant on harvest index in rice under 75% N-** A significant increase in harvest index of 0.38-0.39 was observed in microbial biostimulant conditions 1-3 and HI in control was 0.30. Student’s t-test: significant differences at P < 0.05 and P < 0.01 are represented by ** and ***, respectively.

**Supplementary Fig 7- Soil and plant nutrient analysis-** Influence of microbial biostimulant on soil NPK levels (a) and plant NPK levels (b). Student’s t-test: significant differences at *P* < 0.05 and *P* < 0.01 are represented by * and ** respectively.

**Supplementary Fig 8- Effect of methane derived microbial biostimulant on paddy growth under greenhouse conditions**. Seedlings on the left side represent control and the one on the right side is plants treated with microbial biostimulant. Seedlings 7 days after transplantation

**Supplementary Fig 9-** RT-qPCR based derivative melt curve analysis showing the presence of *M. capsulatus* in paddy roots and leaves.

**Supplementary Table 1-** Indole acetic acid levels observed in microbial biostimulant grown in presence or absence of Tryptophan.

**Supplementary Table 2-** Methane emission reduction from paddy field using methane-derived microbial biostimulant to meet COP26 target for methane reduction by 2030.

**Supplementary Table 3**- List of oligonucleotide primers used in this study.

## AUTHOR CONTRIBUTIONS

SRK, EMD, GJP, KS, KL, CB, GN, PSA, KZ, GR, PS, SA, BRB, PB and CSK performed the experiments. SRK, GJP, EMD, FS, SA, PT and ES analysed the data. SRK, GJP, MUP, VMLK, FS, PT and ES conceived and coordinated the research. SRK, GJP, PT and ES wrote the manuscript.

## CONFLICTS OF INTEREST

The authors declare no conflict of interest.

## DATA AVAILABILITY STATEMENT

All relevant data can be found within the manuscript and its supporting materials.

## Notes

### Competing Interest Statement

The authors have declared no competing interest.

