## Supporting Information for "Enabling Greenhouse Gas Emission Reduction while Improving Rice Yield with a Methane-Derived Microbial Biostimulant"

**1a**

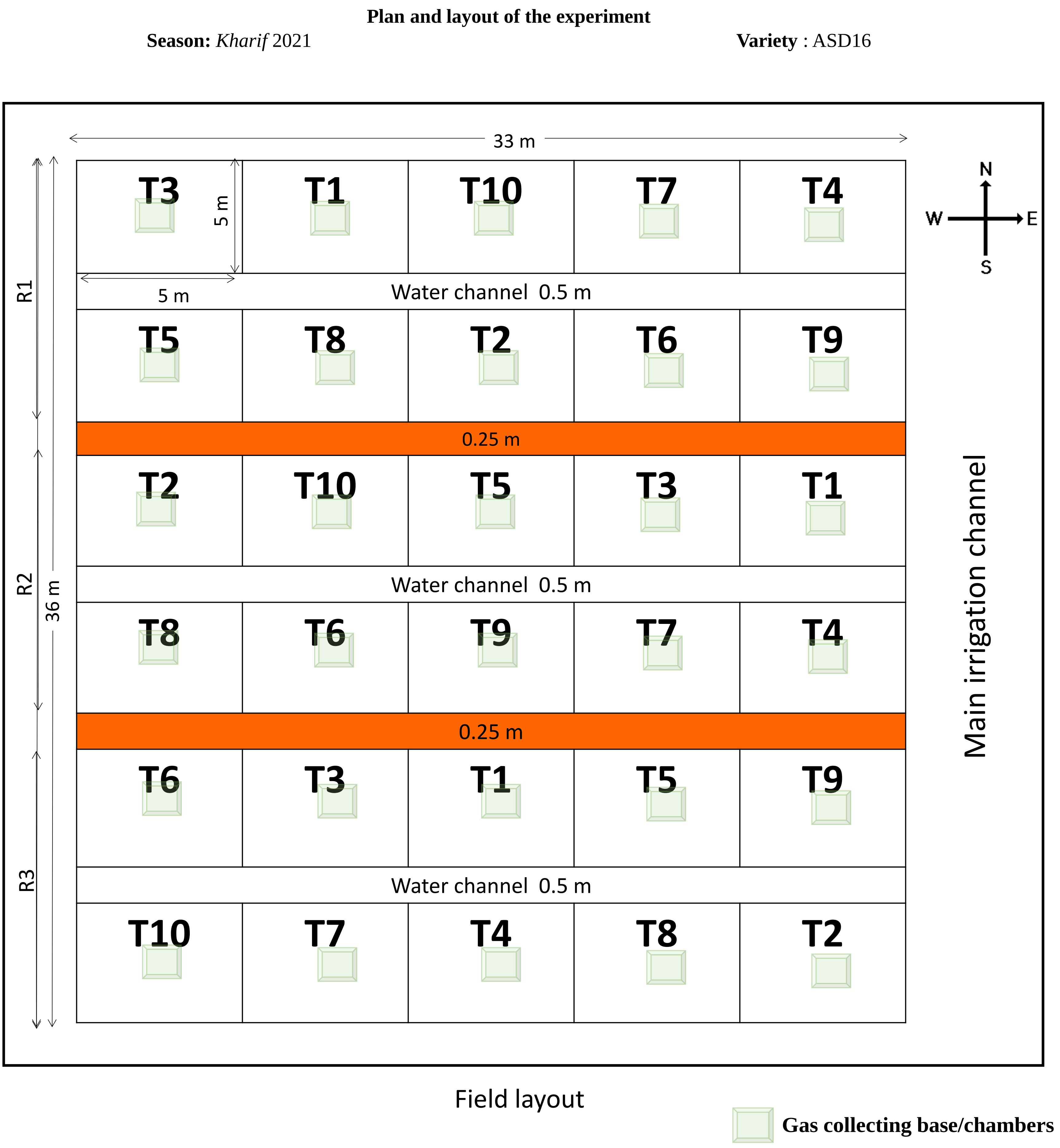


**Supplementary Fig 1a- Experimental field layout for season I testing-** Treatment details are below: T1- Control (100% NPK); T3 & T4- 10 ml/L dose of methane derived microbial biostimulant (100% NPK); T5- Control (75%N); T6- 75%N+microbial biostimulant 5ml/L (condition 1); T7- 75%N+ microbial biostimulant 10ml/L (condition 2); T8- 75%N+ microbial biostimulant 15ml/L (condition 3); T2, T9 and T10 are outside purview of this manuscript and hence are not discussed/explained. R1, R2 and R3 respectively corresponds to replication 1, 2 and 3. Small green box indicate position of gas collection base & chambers.

**1b**

**Plan and layout of the experiment**

**Season:** *Rabi* 2022 **Variety** : ASD16

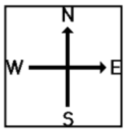


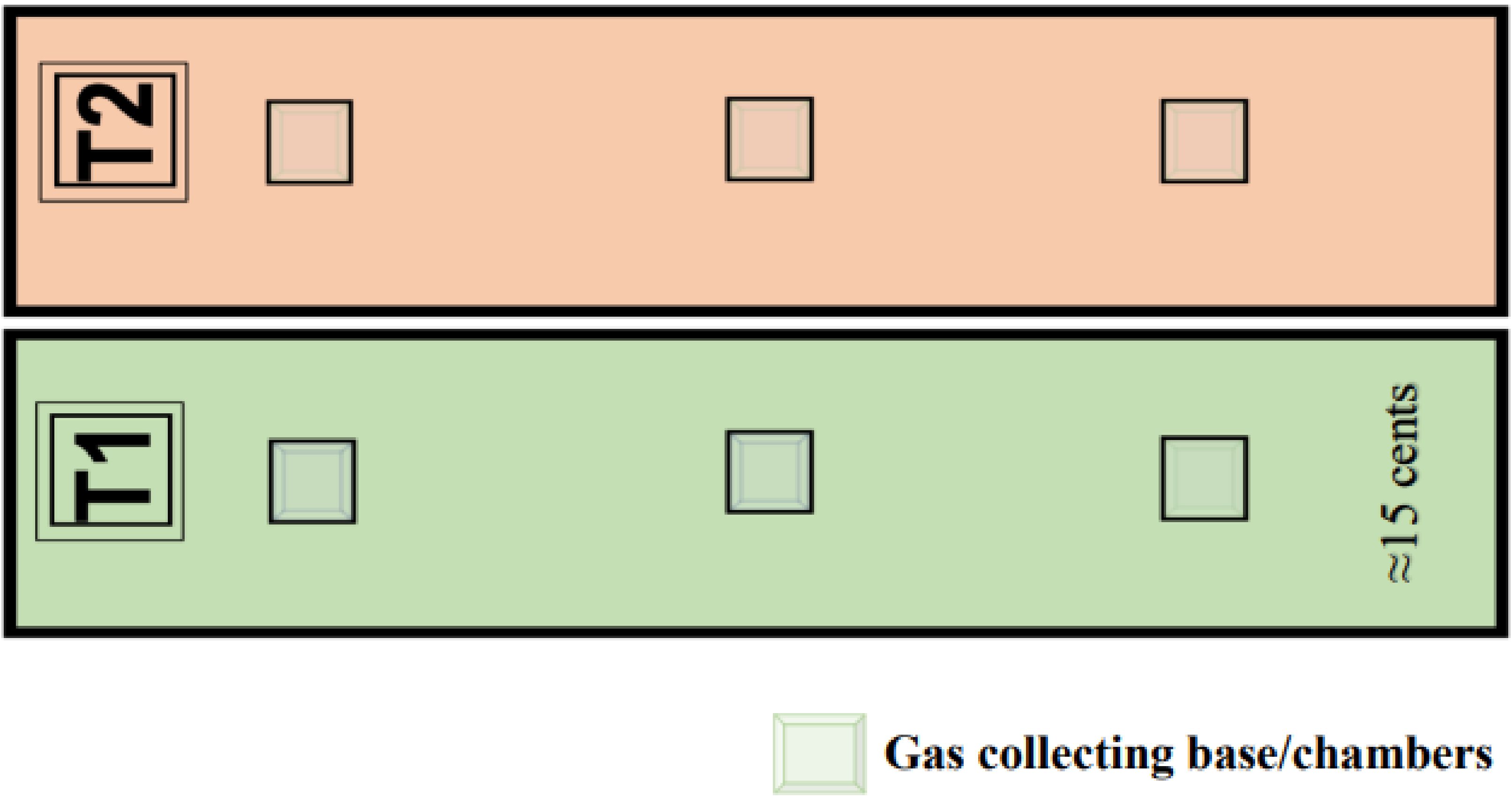


**Supplementary Fig 1b- Experimental field layout for season II testing** - Treatment details are below: T1- Control (100% NPK); T2- Microbial biostimulant- 10ml/L (100%NPK). Small square box indicate position of gas collection base & chambers.

**2a**

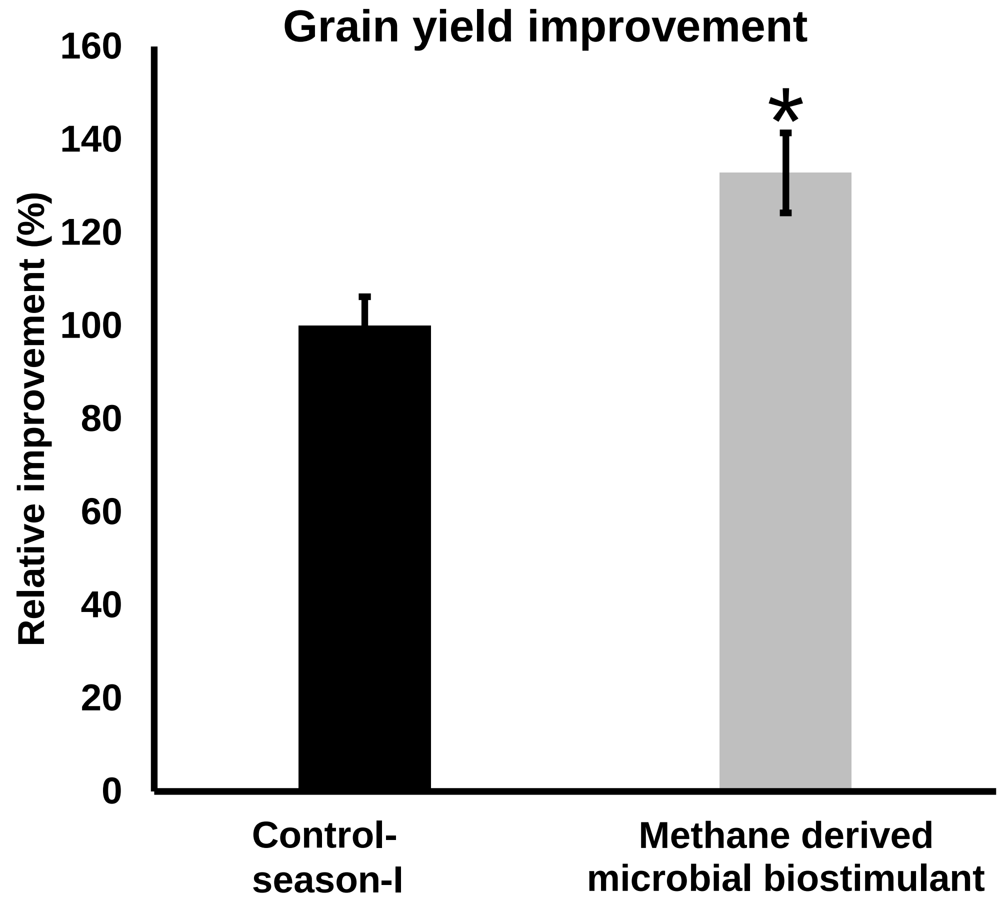


**2b** **2c** **2d**

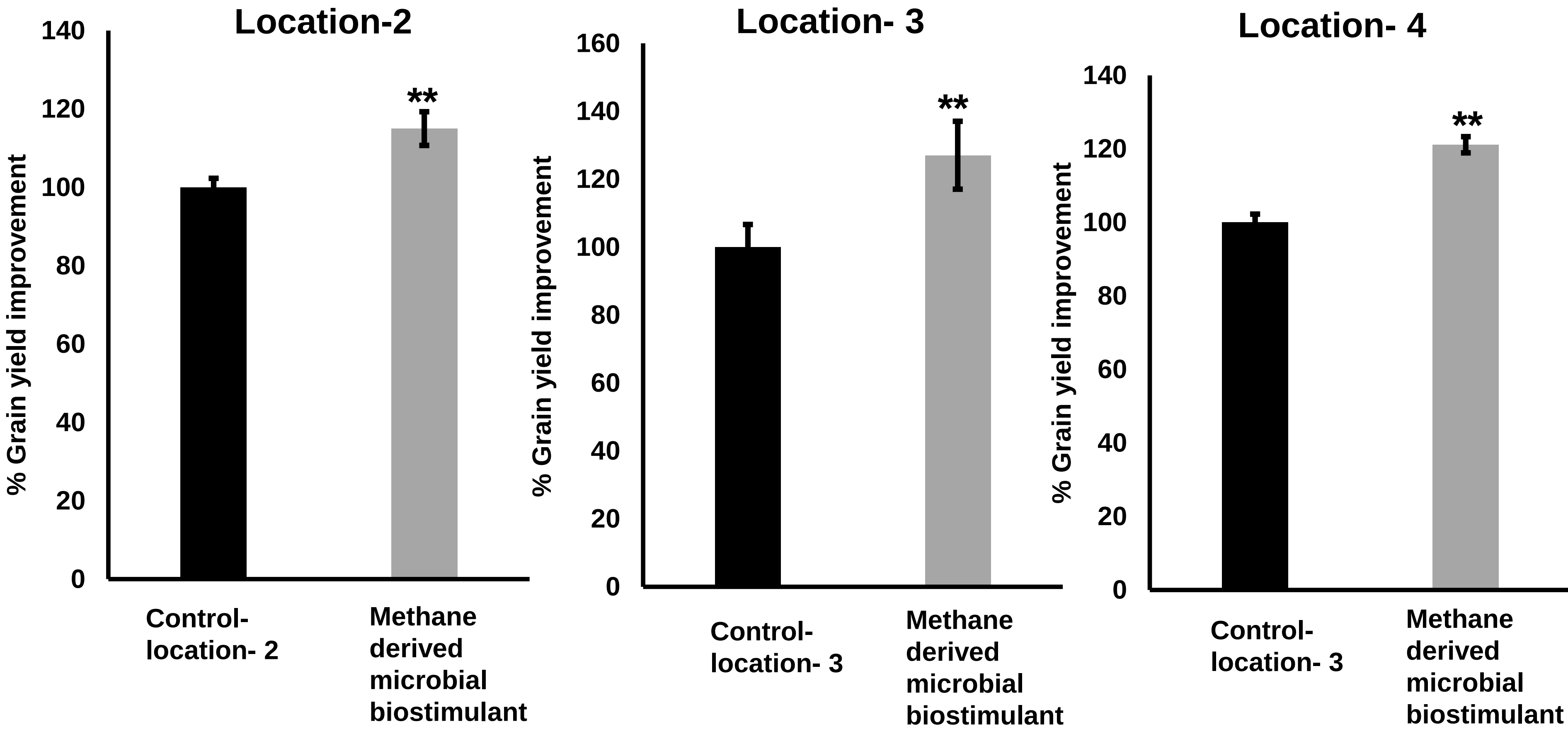


**Supplementary Figure 2a-** Influence of methane derived microbial biostimulant on grain yield where second application was given as foliar spray instead of soil spray.

**Supplementary Figure 2b-d- Multilocation microbial biostimulant validation data-** Grain yield improvement mediated by methane derived microbial biostimulant under different agro ecological locations in India. Differences were evaluated using the two-tailed Student's *t* test and significant differences at $P < 0.05$ and $P < 0.01$ are represented by * and ** respectively.

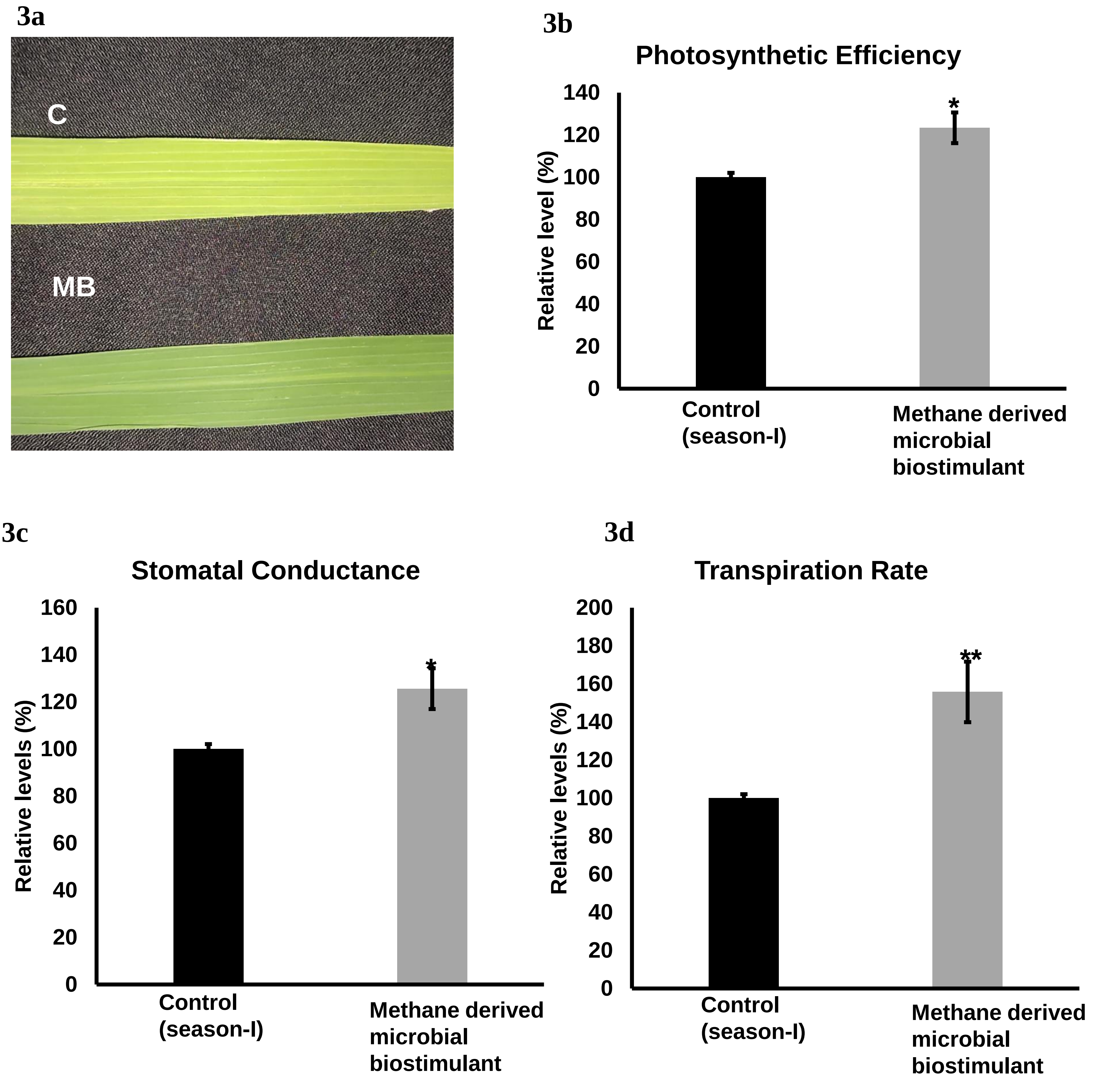


**Supplementary Fig 3(a)- Phenotypic feature of microbial biostimulant treated paddy leaves-** Influence of methane derived microbial biostimulant on greenness in paddy leaf : Control leaf (C) and methane derived microbial biostimulant treated leaf (MB).

**Supplementary 3(b-d)- Effect of methane derived microbial biostimulant on physiological traits in paddy leaves-** Photosynthetic efficiency, stomatal conductance and transpiration rate are represented as % relative to control plants. Student's *t*-test: significant differences at $P < 0.05$ and $P < 0.01$ are represented by * and ** respectively

**4a**

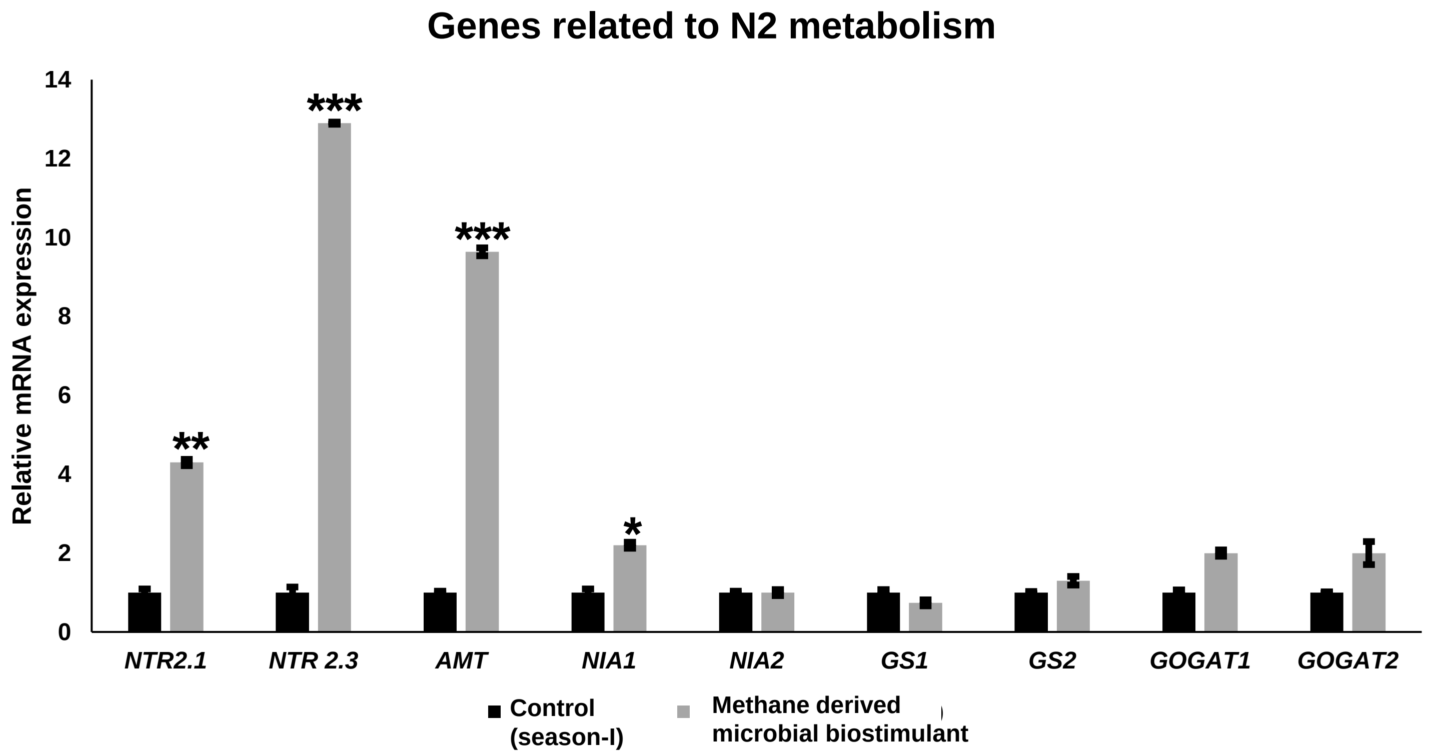


**4b**

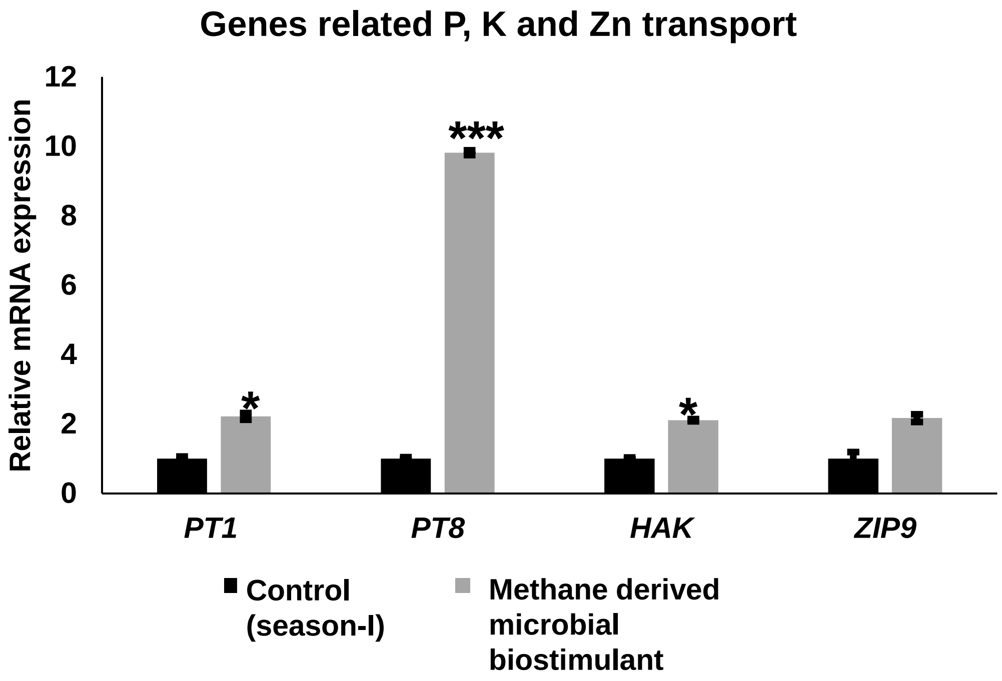


**Supplementary Fig 4- Influence of methane derived microbial biostimulant on expression of root nutrient uptake and transporter genes.**

RT-qPCR analysis showing the expression of genes related to macronutrinet transport and metabolism in roots of microbial biostimulant treated plants. Expression levels of genes were normalized to the endogenous reference gene actin and are represented relative to respective control roots, which was set to 1. Pooled root samples from control and microbial biostimulant treated roots used for RNA extraction. The results shown are from three independent experiments. Error bars indicate mean ± SE. Student's t-test: significant differences at $P < 0.05$, $P < 0.01$ and $P < 0.001$ are represented by *, ** and ***, respectively.

Nitrate transporter (NRT); Ammonium transporter (AMT); nitrate reductase (NIA); Glutamine synthetase (GS); glutamate synthase (GOGAT); Phosphate transporter (PT); High affinity potassium transporter (HAK); Zinc transporter (ZIP).

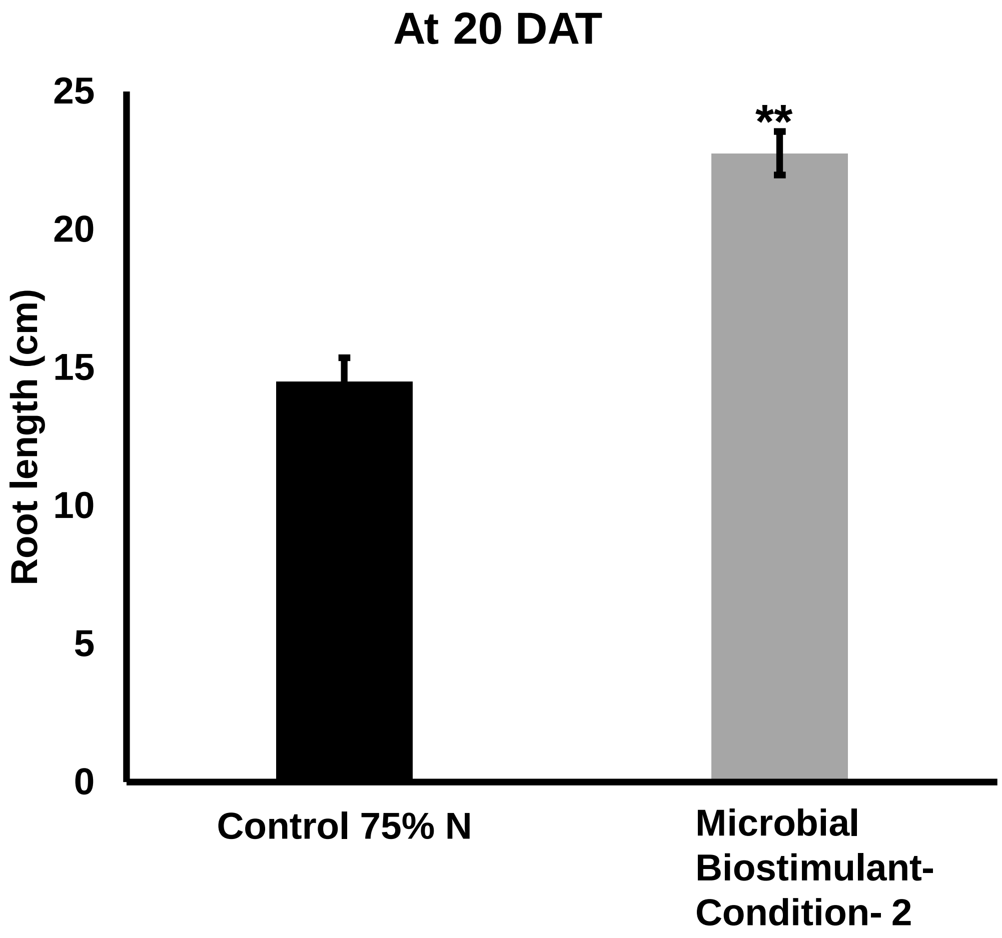


**Supplementary Fig 5- Effect of microbial biostimulant on root length-** Seedling root dipping was performed in paddy roots with microbial biostimulant and twenty days after transplanting. seedlings were uprooted and root length was measured. Student's t-test: significant differences at $P < 0.01$ is represented by "**".

**S6**

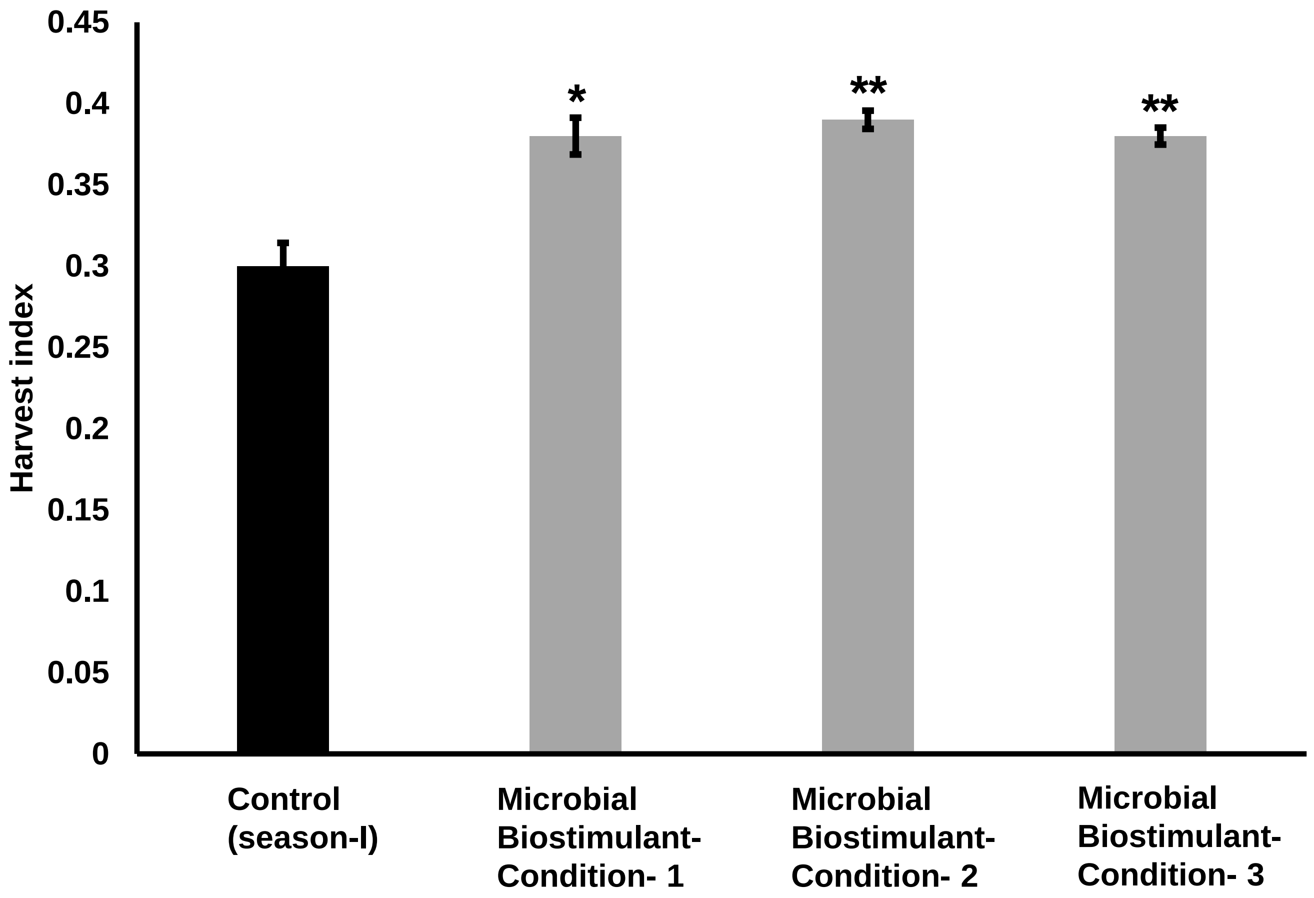


**Supplementary Fig 6- Impact of methane derived microbial biostimulant on harvest index in rice under 75% N-** A significant increase in harvest index of 0.38-0.39 was observed in microbial biostimulant conditions 1-3 and HI in control was 0.30. Student's t-test: significant differences at $P < 0.05$ and $P < 0.01$ are represented by ** and ***, respectively.

**7a**

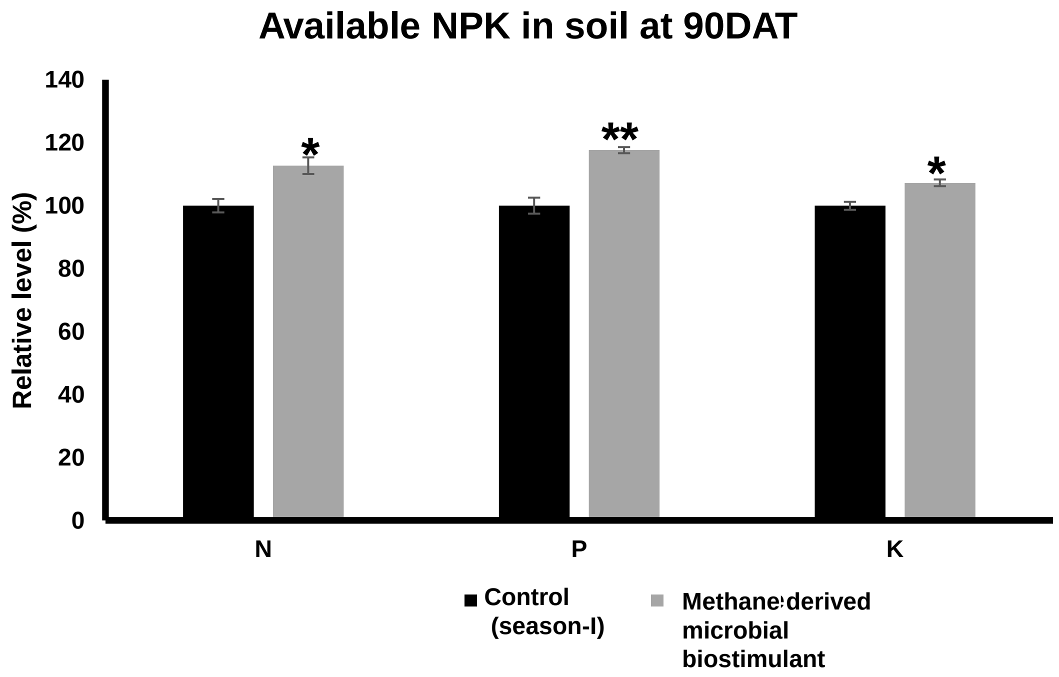


**7b**

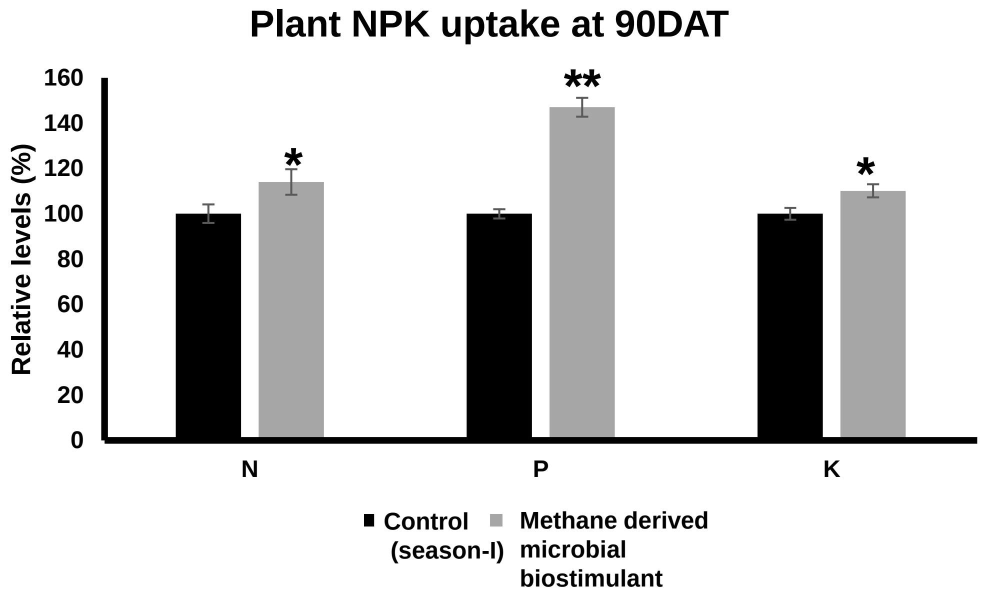


**Supplementary Fig 7- Soil and plant nutrient analysis-** Influence of microbial biostimulant on soil NPK levels (a) and plant NPK levels (b). Student's t-test: significant differences at $P < 0.05$ and $P < 0.01$ are represented by * and ** respectively.

**8**

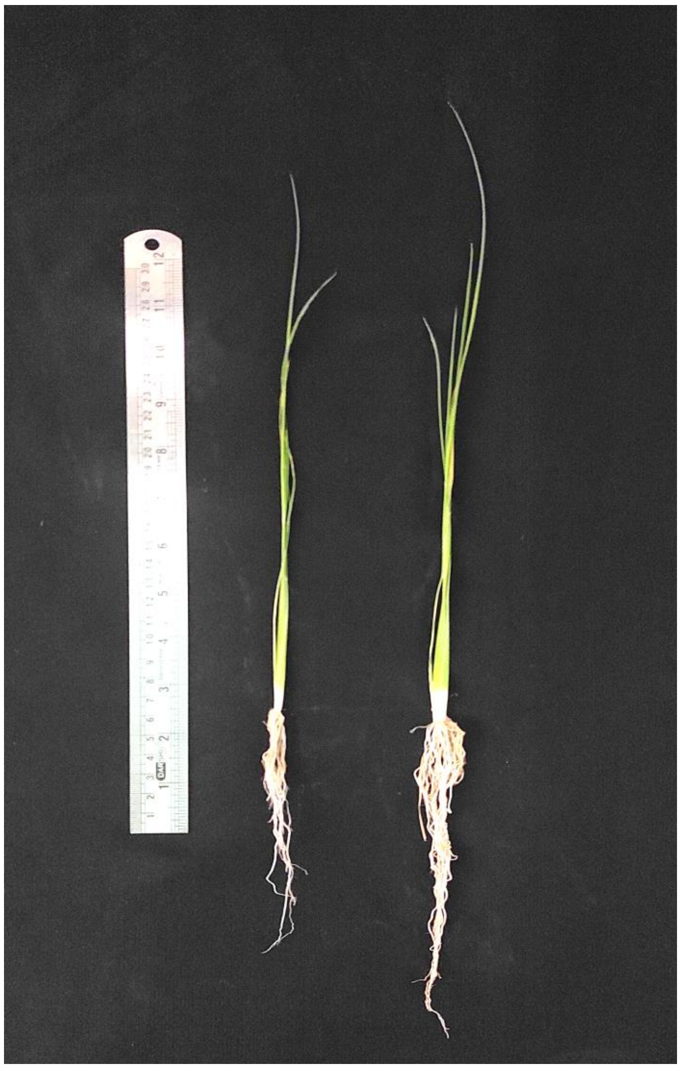

**Supplementary Fig 8- Effect of methane derived microbial biostimulant on paddy growth under greenhouse conditions**. Seedlings on the left side represent control and the one on the right side is plants treated with microbial biostimulant. Seedlings 7 days after transplantation

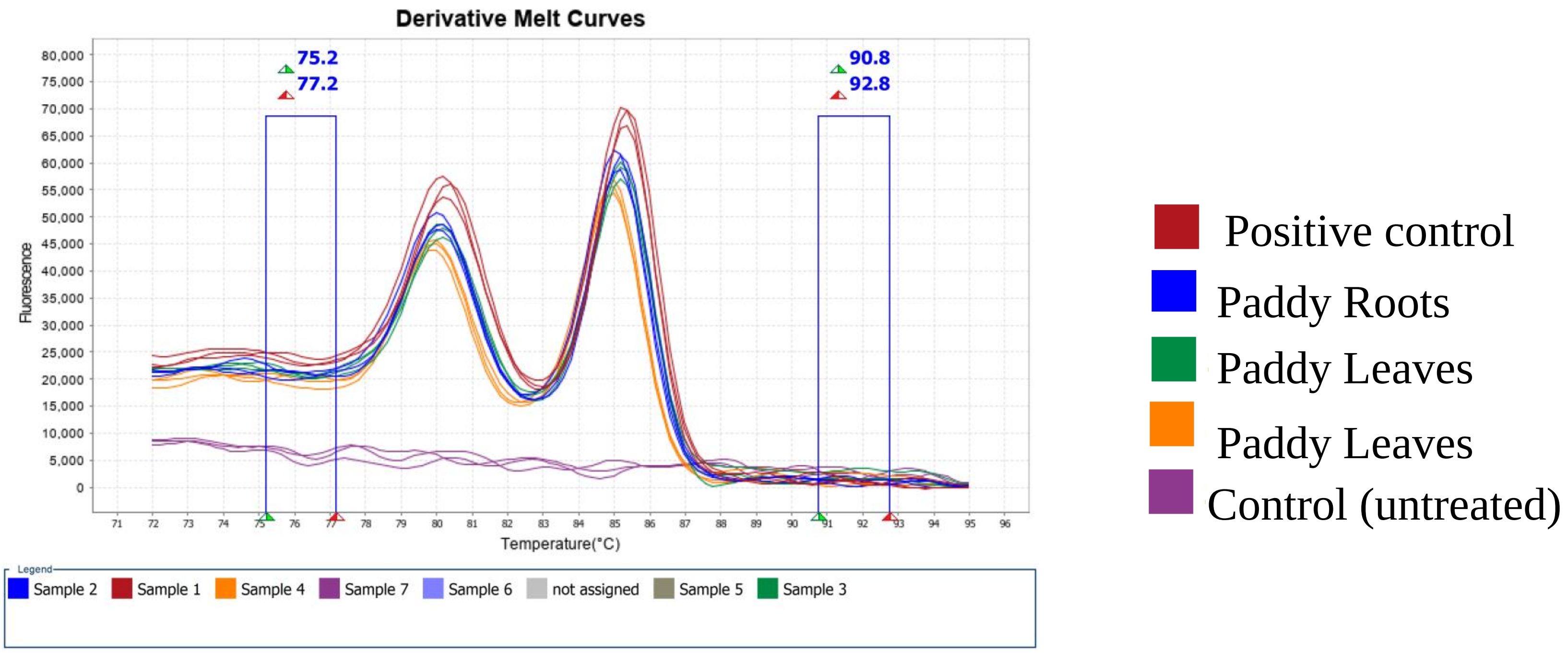


**Supplementary Fig 9-** RT-qPCR based derivative melt curve analysis showing the presence of *M. capsulatus* in paddy roots and leaves.

| Sample | IAA concentration (mg/L) |
|---|---|
| Microbial biostimulant without Tryptophan | 0 |
| Microbial biostimulant with Tryptophan | 1.83-3.61 |

**Supplementary Table 1-** Indole acetic acid levels observed in microbial biostimulant grown in presence or absence of Tryptophan.

| | | | |
|---|---|---|---|
| Methane emission (million metric ton) | 380 | | |
| Methane emission from paddy - 10% contribution (million metric ton) | 30.4 | | |
| Global land under paddy cultivation (million ha) | 162 | | |
| Reduction in methane emission from methane-derived microbial biostimulant (percentage per ha) | 60% | | |
| Global targeted methane reduction per annum (2% of total - COP26 target) million metric tonnes | 7.60 | | |
| Percentage of global paddy cultivation area targeted formethane-derived microbial biostimulant application | 10.00% | 30.00% | 50.00% |
| Actual paddy cultivated area targeted for methane-derived microbial biostimulant application (million ha) | 16.2 | 48.6 | 81 |
| Total methane emission from target area (million metric tonnes) | 3.04 | 9.12 | 15.2 |
| Methane emission from methane-derived microbial biostimulant use in target area (million metric tonnes) | 1.824 | 5.472 | 9.12 |
| Percentage of global methane target achieved | 24% | 72% | 120% |

**Supplementary Table 2-** Methane emission reduction from paddy field using methane-derived microbial biostimulant to meet COP26 target for methane reduction by 2030.

| Gene | Gene Accession No. | Forward Primer sequence (5' - 3') | Reverse Primer sequence (5' - 3') |
|---|---|---|---|
| Os*Actin* | AB047313.1 | ACCATTGGTGCTGAGCGTTT | CGCAGCTTCCATTCCTATGAA |
| *OsPsaH* | Os05g0560000 | GAGGACATCGGCAACACCA | GCCCTTCTTGATCGGAAGCA |
| *OsFD1* | LOC_Os08g01380 | AGCAACAAGCTGGGAGACAG | GAGTAAGGCAGGTCGATCCC |
| *OsET* | LOC_Os04g33630 | GTGGGTGGCGGTGGCAAG | CACGACGGGCCTCAGCTC |
| *OsPsbD* | LOC_Osp1g00170 | CTGCTACTGCTGTTTTCT | GATGTTATGCTCTGCCTG |
| *OsPsbP* | Os07g0141400 | CACGGAGTTCATCGCCTACA | AAGCAAGAAGTCGACCTGGG |
| *OsPsbR3* | Os08g0200300 | GAGGTGGCATGTCAGTCGAT | TTTGCGCCGTACTTGTCAAC |
| *OsPsbS1* | LOC_Os01g64960 | GCTGTTCGGCTTCACCAAGG | ACGCCGGTCTCGATGTTGA |
| *OsCAB1R* | LOC_Os09g17740 | GTTCTCCATGTTCGGCTTCT | GACGAAGTTGGTGGCGTAG |
| *OsCAB2R* | LOC_Os01g41710 | TGTTCTCCATGTTCGGCTTCT | GCTACGGTCCCCACTTCACT |
| *OsCP24* | XM_015781325.2 | CCTTAACCTTAACGGCCATTTCC | CCCGCAGTAAAATCCTGAGC |
| *OsOEP3* | Os07g0544800 | AGCCGCTCATCGAGAAGAAG | GTACTTCTCCGCCTCTTCGG |
| *OsTLP* | LOC_Os08g39430 | GGAGGGAGACGGGGTGGG | TCAGAAAGGGAGGAGAGCCGT |
| *OsLHC2.1* | LOC_Os02g52650 | GGCGCGGTCGAACGAGCT | GTTTAGCCATTAGTCAAAGCAATC |
| *OsChlI* | LOC_Os03g36540 | CTTCACCGTCTGCAATGTAG | GATCTTAGGGTCGATGACGTT |
| *OsChlH* | LOC_Os03g20700 | GCACGGGAACTTGGCGTTTCATTA | ACATGTCCTGGAGCTGCTTCTCAT |
| *OsChlD* | LOC_Os03g59640 | TAGCACAGCTGTCAGAGTGGGTTT | TTGCCAGCCACCTCAAGTATCTCA |
| *OsHEMA* | AB011416.1 | GATGCAATCACTGCTGGAAAGCGT | CCATCTTGCCAGCACCAATCAACA |
| *OsHEMC* | LOC_Os02g07230 | GAGCTGTCTCATTCAGATCTGT | AGGGATTACAATAACCGTGGAA |
| *OsYGL13* | LOC_Os08g06630 | CTCTCCCACAAGATCAAGGATG | GTTTGTATGAGGGCTCATTTCC |
| *OsYGL8* | Loc_Os01g73450, | TGGATCTAACATGACACGCACCCA | ACTGTAACGGCATTCTTCTCCGGT |
| *OsRCA* | U74321.1 | CTCTTCGTGCCCGTGTTTAC | TCGGAGTTAGCGTCACCAAG |
| *OsRbcS2* | L22155.1 | AGTCTGGTGGCAACTAAGCC | GCACGGCCGGTAAAATCAAA |
| *OsRbcS3* | AK068555.1 | ACCATCTCAATGGCCTCTGC | TGTGTGCATATAGCCGGAGC |
| *OsRbcS4* | AK070257.1 | AACGTTAGGCAGGTGCAGTT | TGCAGCTTAACACGGACACA |
| *OsRbcS5* | AK099574.1 | GGAGTCCGGCGGAAACTAAG | GGAAACCAATGCAAGGTGGC |
| *OsHYR* | Os03g02650 | CCGAGGGCTTGATGATGAG | CGTAAGCCCATTTCAGGAATG |
| *OsMOC1* | AY242058 | CTGCTCCGGCTGCACTAC | CCACGCTGAGACGGAGAG |
| *OsSLR1* | Os03g0707600 | GAGTCGCTGCACTACTACTCC | GCGCGTGTGCCAGCCCAGCGT |
| *OsLAX2* | AB669025 | TTCTTGCCTCAGATTCCAAG | CCTCTGCATGTTATCTCCAC |
| *OsMADS57* | LOC_Os02g49840 | CAGATTATGTTGTCGGATGCTC | AAAGCAATAGAGAGTAAGCAGGGT |
| *OsHSF2AD* | LOC_Os03g06630 | CAGCAGGCACTTGGCACC | TTCTTGTCACGCTTTAGCCTGT |
| *OsCKX11* | Os08g0460600 | CAACGCAATCATTGACGCC | TTGCACCCCTCCCAAATGT |
| *OsRGN1* | LOC_Os01g49160 | CGGCTACACCGACCAGGAG | CGCGATGATGGACCACCT |
| *OsNOG1* | MF687920 | TCCGACTTACAATGAACAC | GGTAGCAGGACTCCACTT |
| *OsSPL9* | LOC_Os05g33810 | AGATGGGCAGGTGATTAT | TGTGGGAGAGCTTTAGTC |
| *OsIPA1* | LOC_Os08g39890 | CGGTCGACTAGCTGCATCTGTTGG | CATCGTGTTGCTGGTTTGGTCGAAG |
| *OsSPL14* | Os08g0509600 | CAAGGGTTCCAAGCAGCGTAA | TGCACCTCATCAAGTGAGAC |
| *OsLC2* | AK101341 | AGCATCAGCTTTGGACGAGGA | CAGTTGGTGGAATAGAGCCAGAAT |
| *OsVIL2* | XM_015764905.2 | GGAGTATGCTTTCCGGATCA | GTGGGAAACAACATGTGCAG |
| *OsGRF4* | LC333011.1 | GAAAGCCTGTGGAAACGCA | CAACGCCGAGCCAAATGAG |
| *OsNRT2.1* | NM_001401744.1 | CTTCACGTCGTCGAGGTACT | CACTCGGAGCCGTAGTAGTG |
| *OsNRT2.3A* | XM_015773038.2 | CGCTGCTGCCGCTCATCCG | CCGTGCCCATGGCCAGAC |
| *OsNIA1* | XM_015767224.2 | TCAAGGTGTGGTACGTGGTG | CGAGGTCATAGCCCATCTTC |
| *OsNIA2* | NM_001422594.1 | TGTACCAGGTCATCCAGTCG | CGATGACGTACCACACCTTG |
| *OsAMT1* | AF001505.2 | GGTTTCTCTCCCTCTCCGAT | CCACCTTCACACCACACATT |
| *OsGS1* | AB180688.1 | TGTTTCTCCTCATCCCTGC | TCACAGTCCTCGCTTTGC |
| *OsGS2* | X14246.1 | GGAGAGGTCATGCCTGGTCAGT | ACTACACCAGCCTGCTCCGTTA |
| *OsGOGAT1* | XM_015793761.2 | GTGCAGCCTGTTGCAGCATAAA | CGGCATTTCACCATGCAAATC |
| *OsGOGAT2* | NM_001402625. | CCTGTCGAAGGATGATGAAGGTGAAACC | TGCATGGCCCTACTATCTTCGCATCA |
| *OsPT1* | AF536961.1 | CGCTTCCGTACGAGTGGTAGT | GGTTCTTTCAAATCCAGGGAAA |
| *OsPT8* | AF536968.1 | AGAAGGCAAAAGAAATGTGTGTTAAAT | AAAATGTATTCGTGCCAAATTGCT |
| *OsHAK1* | NM_001402365.1 | GTTGATGATGCTGATGTTGGAAG | CCAACACTTTCAGCTGAAAC |
| *OsZIP9* | NM_001420533.1 | CATCAGTTCTTCGAAGGGATAGG | TGTGGTTAGCGAGAAGAAGATG |
| *McMopB* | AF031148.1 | ACAGGCCGAAGAGACTTTCA | GGTGGTGTTGCCTTCGTA AT |

**Supplementary Table 3-** List of oligonucleotide primers used in this study.
