## Supporting Methods for "Enabling Greenhouse Gas Emission Reduction while Improving Rice Yield with a Methane-Derived Microbial Biostimulant"

#### Field trial design for validation of microbial biostimulant

The season I study was conducted from July to October 2021 at Lat.12°58'05.56"N, Lon.079°09'39.96"E at VIT School of Agricultural Innovations and Advanced Learning, Vellore Institute of Technology, Vellore, Tamil Nadu, India. The trial covered a total area of 29 cents, with 30 plots prepared in dimensions of 5m x 5m. The experimental design followed the Randomized Block Design (RBD) method, with ten treatments and three replicates. T9 and T10 are outside purview of this manuscript and hence are not discussed/explained. Season II study was carried out at between February to June 2022 at Lat.12°58'45.1"N, Lon.79°09'47.9"E in a farmer's field near Vellaikalmedu, Vellore, Tamil Nadu, India. The testing area covered an area of 810 m2 for each treatment. In both cases, recommended nitrogen doses were made as split applications: at basal, tillering, and panicle initiation stages. Recommended dose of fertilizers i.e. 40: 20: 20 was applied NPK/acre was applied. The fertilizer was supplied through different sources like urea (46% N), single super phosphate (16% $P_2O_5$) and muriate of potash (60% $K_2O$) as per the package of practice. For treatments with only 75% of N, half dose of N, full dose of $P_2O_5$ and $K_2O$ were applied as basal dose. Remaining N was split into two doses at tillering stage and at booting stage of the crop. The field was carefully prepared for the trials with bunds and buffer channels in place to prevent cross-infiltration and separate each plot. Each plot was allocated a specific irrigation channel to ensure that water applied to one plot remained contained within that plot. This design effectively prevented cross-infiltration. To maintain uniform plant population, during transplantation, 2-3 seedlings were planted per hill with a spacing of 20cm X 10cm, resulting in 50 hills per $m^2$. Variety used for season I and season II trials are ASD-16 released by Tamil Nadu Agricultural University, Coimbatore, India.

#### $CH_4$ derived microbial biostimulants application

Roots of twenty days old seedlings were immersed in microbial biostimulant solution for 20 minutes, prior to transplantation to the main field. A soil (for T3 during season I) or foliar spray (for T4 during season I) during peak tillering stage and foliar spray at panicle development stage were also performed as second and third applications respectively.

### Multilocation Trials

To understand the efficacy of microbial biostimulants on improving grain yield under different agro-ecological regions, multilocation trial was conducted at Jhansi, Uttar Pradesh, India (GPS coordinate: 25.5101° N, 78.5411° E), Jabalpur, Madhya Pradesh, India (GPS coordinate: 23.2152° N, 79.9601° E) and Mandya, Karnataka, India (GPS coordinate: 12.5690° N, 76.8107° E). Crop management practices recommended for each location was followed. Seed variety used were Pusa Basmati 1121 (Jhansi, India), JR-206 (Jabalpur, India) and MTU-1001 (Mandya, India). Microbial biostimulant applications at the rate of 10ml/L were performed at three times during the crop growth. Control plots received water sprays.

### Analysis of physiological parameters in paddy

Physiological parameters were analyzed from five tagged plants from each treatment during peak vegetative growth stage. Photosynthetic rate was analysed using Infrared gas Analyser (IRGA, Licor-6800; Li-Cor inc., Lincoln, NE, USA).  Leaf stomatal conductance and transpiration rate was analysed using a LICOR 6800 portable photosynthesis system (Lincoln, Nebraska, USA). These observations were recorded on clear sunny days between 10:00 am and 12:00 noon with a saturated light environment.

### RNA extraction and Reverse transcriptase- quantitative polymerase chain reaction

RNA from leaves (collected during peak tillering stage) and panicles (collected during panicle initiation stage) from control and treatments were isolated as reported previously (Kumar et al., 2018). Leaf and panicle RNA were normalized using *Osactin* that was previously reported as an appropriate endogenous gene in rice. Fold-change differences in gene expression were analyzed using the comparative cycle threshold (*Ct*) method. Relative quantification was carried out by calculating *Ct* to determine the fold difference in gene expression [$\Delta Ct$ target – $\Delta Ct$ calibrator]. The relative level was determined as $2^{-\Delta\Delta Ct}$. Primers used for the analysis are mentioned in **Supplementary table 3**.

### Isolation and confirmation by microbial cells in rice roots and leaves by quantitative polymerase chain reaction

The presence of *M. capsulatus* in rice roots were evaluated by seedling root dip treatment and foliar application followed by re-isolation. Roots of paddy seedlings (25 days old) were dipped in microbial biostimulant suspension for 20 min. For control plants, the roots were dipped into water. Treated seedlings were transplanted in pot containing soil + farmyard mannure (1:1). Another set of plants were sprayed at 10ml/L dose of microbial biostimulants whereas control plants received water spray. Plants were maintained at the greenhouse facility. One week after transplantation, the seedlings were carefully uprooted, and roots were gently washed with running tap water to remove all soil particles. Root and leaves tissues were selected for isolation and root/leaves samples were surface sterilized by 70% ethanol for 2 min, and then samples were washed thrice with sterile water. Surface sterilized roots/leaves were cut into small pieces or segments of 0.5 to 1cm in size aseptically using a sterile blade. Root segments of 15 to 20 numbers were suspended in conical flask containing 20ml of nitrate mineral salts (NMS) media. The flasks were sealed with suba-seal and head spaces were filled with methane (0.2 bar), incubated @ 45° C at 180rpm in an orbital shaker. At 4$^{th}$ day of incubation, 100µL of cultures were transferred to 20 ml of fresh NMS media with methane as carbon source and continued incubation to enrich the microbial culture. The enriched cultures were used to RT-qPCR to reconfirm the culture identity.

Genomic DNA isolated from the cultures of control and microbial biostimulant treated paddy samples along with positive control (pure genomic DNA isolated from bacterial strain) were subjected for strain specific RT-qPCR using *MopB*- primers followed by High resolution melt curve analysis developed in house. All PCR assays were performed as single-tube assays in triplicates in 0.1-mL strip tubes and caps using Quant studio 3™ system (Applied Biosystems Inc, USA). Each 10 µL reaction mixture contained 0.5 µL of both forward and reverse primers (10pM), DNA template (100ng), 5µL of Melt doctor reagent™ (Applied Biosystems; Catalog number:4425557) and 3.5 µL of MilliQ water. Each sample was incubated for five minutes at 95 °C, then was denatured for 10 seconds at 94 °C, annealed for 10 seconds at 60 °C, extended for 10 seconds at 72 °C for 40 cycles. A post-PCR HRM analysis was performed from 72 °C to 95 °C, increasing at 0.2 °C/step. The results were analyzed using high resolution melt software (Applied Biosystems Inc, USA).

**Extraction and quantification of IAA**

The IAA production by the *M. capsulatus* was quantified by growing cells for 3 days at 45 °C in Nitrate Mineral Salt (NMS) broth supplemented with and without 5mM tryptophan. After incubation, the broth was centrifuged at 5000×g for 10 min, and IAA was quantified in the supernatant by high performance liquid chromatography (HPLC). The instrument (Model:

1260 Infinity II with quaternary pump, autosampler & VWD detector, Agilent Technologies) equipped with $C_{18}$ symmetric reverse phase column (Poreshell 120- 4.6 × 250 mm, 4 µm, Agilent Technologies) was used for analysis. Separation was carried out in isocratic mode with mobile phase consisting of 0.1% acetic acid (A) and Acetonitrile (B) at a ratio of 40:60 A:B. The column temperature was maintained at 30°C with a flow rate of 0.5 ml/min. The total run time was 15 minutes. Data was extracted at 287 nm and peak area obtained from standards and samples were used to quantify the levels of IAA.

#### Soil and Plant Nutrient Analysis

The plant, grain and soil analysis methods were carried out as per Motsara and Roy (2008). A random sampling consisting of fiver plants per plot was taken for plant and grain nutrient analysis. Plant/grain samples (minimum from five tagged plants) were collected from each treatment at 90 DAT and they were cleaned and shade dried. Later the shade-dried samples were oven-dried at 60 ± 5º C for 24-48 hours. The samples were finely powdered using mixer grinder. The finely ground plant samples were used for analysis. Total nitrogen was estimated using micro Kjeldahl digestion and distillation method (Jackson, 1973). Digestion of plant samples were carried out with di-acid mixture. Exactly 0.5 g of powdered plant sample was pre-digested with 5 ml of concentrated $HNO_3$ and digested with a di-acid mixture (HNO3:HClO4 in the proportion of 9:4 ratio). Digested samples were diluted with distilled water and volume was made up to 100 ml. The same samples were preserved for P and K analysis. To determine total phosphorus, vanadomolybdophosphoric yellow colour method was used and the phosphorus content in the digest was estimated using spectrophotometer (Tandon, 2005). Total potassium content in the digested samples was estimated using flame photometer (Tandon, 2005). To analyse micronutrients, 0.5g of 100 mm mesh powdered sample was digested with diacid (HNO3:HClO4 in 9:4) mixture in a digestion chamber. After complete digestion, it was filtered (Whatman No. 42) to a 25 ml volumetric flask and volume was made by thoroughly washing with deionized water. This sample was preserved for micronutrient estimation by using atomic absorption spectrophotometer.

The uptake of nutrients was estimated using below formula:

$$\text{Uptake (kg ha}^{-1}\text{)} = \frac{\text{Nutrient concentration (\%)} \times \text{Weight of dry matter (kg ha}^{-1}\text{)}}{100}$$

Soil analysis before and after trial was carried out to get the levels of initial and final macronutrient availability. For soil macronutrient analysis, soil samples from all the treatments were taken at 90 DAT. Samples were obtained from the surface (0 to 15 cm). The samples collected were shade dried, finely powdered and sieved using 2 mm sieve. Sieved soil sample was used for analysis. Available nitrogen was determined by modified alkaline potassium permanganate method (Sahrawat and Burford, 1982). To check the levels of available phosphorus, sample was extracted using 0.5 M sodium bicarbonate at pH 8.5 (Olsen et al., 1954). The intensity of colour developed by stannous chloride was measured in spectrophotometer at 660 nm. To determine the levels of available K, sample was extracted with neutral 1N ammonium acetate extract (Hanway and Hiedal, 1952) and the content was determined by flame photometer. The amount of nutrients is expressed as relative percentage to the controls.
